# Prospective and retrospective representations of saccadic movements in primate prefrontal cortex

**DOI:** 10.1101/2022.09.26.509463

**Authors:** Ioana Calangiu, Sepp Kollmorgen, John Reppas, Valerio Mante

## Abstract

Dorso-lateral prefrontal cortex (dlPFC) in primates plays a key role in the acquisition and execution of flexible, goal-directed behaviors. Recordings in monkey dlPFC have revealed possible neural correlates of the underlying cognitive processes like attention, planning, or decision-making, both at the single-neuron and population levels. Integrating these observations into a coherent picture of dlPFC function is challenging, as these studies typically focused on neural activity in relation to a few, specific events within a single, fully learned behavioral task. Here we obtain a more comprehensive description of dlPFC activity from a large dataset of population recordings in monkeys across a variety of behavioral contexts. We characterized neural activity in relation to saccades that monkeys made freely, or at different stages of training in multiple tasks involving instructed saccades, perceptual discriminations, and reward-based decisions. Across all contexts, we observed reliable and strong modulations of neural activity in relation to a retrospective representation of the most recent saccadic movement. Prospective, planning-like activity was instead limited to task-related, delayed saccades that were directly eligible for a reward. The link between prospective and retrospective representations was highly structured, potentially reflecting a hard-wired feature of saccade responses in these areas. Only prospective representations were modulated by the recent behavioral history, but neither representations were modulated by learning occurring over days, despite obvious concurrent behavioral changes. Dorso-lateral PFC thus combines tightly linked flexible and rigid representations, with a dominant contribution from retrospective signals maintaining the memory of past actions.

## Introduction

Dorso-lateral prefrontal cortex (dlPFC) in primates is thought to play a key role in goal-directed behavior by flexibly maintaining and integrating signals required to select contextually-relevant actions, through processes like working memory, attention, and the context-dependent accumulation of sensory evidence^1–7^. This view of dlPFC function has been shaped in particular by studies in primates engaged in saccade-based tasks, many of which focused on characterizing responses preceding an action^8–10^. A large literature on pre-saccadic responses revealed neural dynamics that is strongly context-dependent^9,11^, can support abstract representations^7,12–16^, and reflects representations of task variables that are randomly mixed at the level of single neurons^7,17^, consistent with a primary role of dlPFC in the prospective control of flexible decisions.

Prominent task-related activity, however, has often been reported also during^11^ and following saccades^8,11,18–28^. One widely reported signal is post-saccadic activity, which in several areas of dlPFC is intermingled with pre-saccadic and movement related activity^8,11,18,19^. The proposed functions of post-saccadic activity mostly differ from those of pre-saccadic activity, and include the retrospective monitoring of behavioral context^20,21,27,29^, terminating cognitive processes that select contextually-relevant actions^30^, updating retinotopic maps to ensure visual stability^19,26,31^ or alternatively, the preparation for future actions^11^. Currently, a systematic comparison of the prevalence and properties of pre- and post-saccadic activity across contexts, stages of learning, and neurons is lacking, and consequently the primary function of dlPFC remains a matter of debate.

Here, we compared neural population recordings obtained with chronically implanted electrode arrays in dlPFC of macaques^32,33^ across a variety of behavioral contexts. Such array recordings arguably provide a more unbiased view onto the signals represented by a neural population compared to past single-neuron recordings. Monkeys were engaged in several classic, saccade-based, motor and decision-making tasks, which however differed in a critical aspect from past studies^9,11^. Operant saccades were not only preceded, but also followed, by a delay period that was randomized from trial-to-trial^20^, simplifying a direct comparison between the prevalence and properties of pre-saccadic and post-saccadic representations. To establish the role of behavioral context on the inferred dlPFC representations, we compared neural population responses across tasks, across different stages of learning, and between trained and freely chosen saccades.

We find that dlPFC can represent saccade direction from the time of planning, through movement, until the resulting outcome and beyond. Notably, the dominant signal across tasks, saccade types, and learning is post-saccadic activity, suggesting a key role of dlPFC in retrospective computations. Our findings are organized in three sections. First, we show that post-saccadic activity overall is stronger than, and distinct from, pre-saccadic activity (Fig. 1-4). Like pre-saccadic activity, post-saccadic activity is persistent and inherently tuned to the direction of the saccade, but it represents the past rather than the future action^8,11,18,19,21–28^. Unlike pre-saccadic activity, post-saccadic activity appears to occur in relation to every saccade, albeit with some modulation due to the behavioral context. Second, we show that some components of the identified saccadic representations have tightly linked pre- and post-saccadic dynamics at the single neuron and population level (Fig. 5-6), consistent with a “hard-wired” feature of the underlying circuits. Third, we study how the representations of saccade direction are shaped by learning on short^34^ (consecutive trials) and long time-scales^35^ (days and months) in an associative-learning task^24,34^(Fig. 7-8). Only pre-saccadic representations are influenced by recent trial history, and both pre- and post-saccadic representations show little or no modulation on the longer time-scales associated with large changes in behavior. Overall, these findings imply that rigid, structured representations are a key component of dlPFC computations, with a dominant contribution from post-saccadic signals maintaining the memory of past actions.

**Figure 1.**
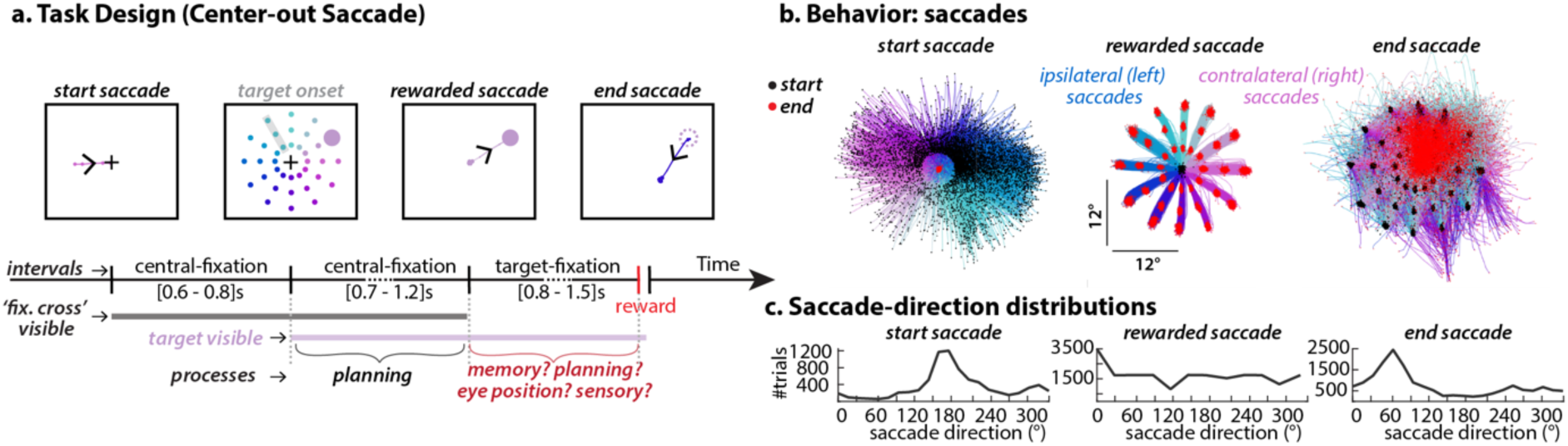
Instructed-saccade task and behavior. **a.** Subjects performed a visually-guided, delayed-saccade task. A trial was initiated by a saccade to the fixation point after which monkeys were required to maintain fixation (“central-fixation”). After a randomized delay a saccade target was presented in the periphery (in total 33 unique positions per experiment for monkey T and 24 for monkey V and C, Supp. Fig. 1c). Here we show 36 unique positions, pooled across all experiments of monkey T and highlight a direction (3 positions) that was not presented in a particular experiment. See Methods for a full description of how targets were shown in each experiment. When the fixation point disappeared (a second randomized delay), monkeys were required to execute a saccade to the target. After the saccade, monkeys were again required to maintain fixation, this time on the target, for the duration of a final random interval (“target-fixation”). **b.** Top: Eye-trajectories for three types of saccades: **start** saccades (to the fixation point) that initiate the trial; **rewarded** saccades (to the visual target); and **end** saccades (from the visual target) with no task constraints. Eye trajectories are sorted by saccade direction. The direction of start and end saccades is discretized into classes that match the experimentally-set directions of the rewarded-saccade in the corresponding session. **c.** Distribution of saccade-direction for the three different saccade types pooled over all sessions and radii. In approx. 45% of trials, the monkey was already at fixation point when the new trial started, thus there are fewer start saccades than end saccades. Moreover, we only analyze start and end saccades with amplitudes similar to the experimentally-set amplitudes of the rewarded saccade, i.e. between 4 and 16 deg.

## Results

### Behavioral task and neural recordings (Fig. 1)

We first consider recordings from three monkeys that were engaged in a visually-guided, instructed-saccade task, requiring them to perform a sequence of saccades and fixations on each trial to obtain a reward (Fig. 1a). We analyzed neural activity and eye movements for all trial epochs (Fig. 1a) and different types of saccades, i.e. the instructed and freely initiated saccades occurring before, during, and after each trial (Fig. 1b). We refer to the initial saccade to the fixation point as the “*start saccade”*, the saccade to the target as the “*rewarded saccade”*, and the first saccade away from the target after reward delivery as the “*end saccade”.* Figure 1c shows the distribution of saccade directions for the different saccade types pooled over all experiments and radii (Monkey T). The start saccade is followed by the “*central-fixation”,* i.e. the initial fixation on the fixation point lasting for a randomized interval (1.3-2s) and preceding the “*rewarded saccade*”. Crucially, the *rewarded saccade* is followed by the “*target-fixation*” lasting for a randomized interval (0.8-1.5s), i.e. the fixation on the target until it disappears (Suppl. Fig. 1b). The inclusion of this prolonged *target-fixation* is a key difference from instructed-saccade tasks used in past studies^8,11,18,19,36–39^ and greatly simplifies the interpretation of post-saccadic neural activity.

Neural activity was recorded with 96-channel Utah-arrays implanted in pre-arcuate cortex, a region of dorso-lateral PFC close to, and possibly including, the most rostral part of the frontal eye fields^40^ (Supp. Fig. 1a). Monkey T and V had the array placed in the concavity of the arcuate sulcus in the left hemisphere, while monkey C had it placed in the right hemisphere, above the principal sulcus^33^. We show results from monkey T and monkey V in the main text. We show results from monkey C in the supplementary figures to highlight similarities and differences from the other monkeys, which may be due to a different array placement. During the duration of an experiment, monkeys were head-fixed.

**Figure 2.**
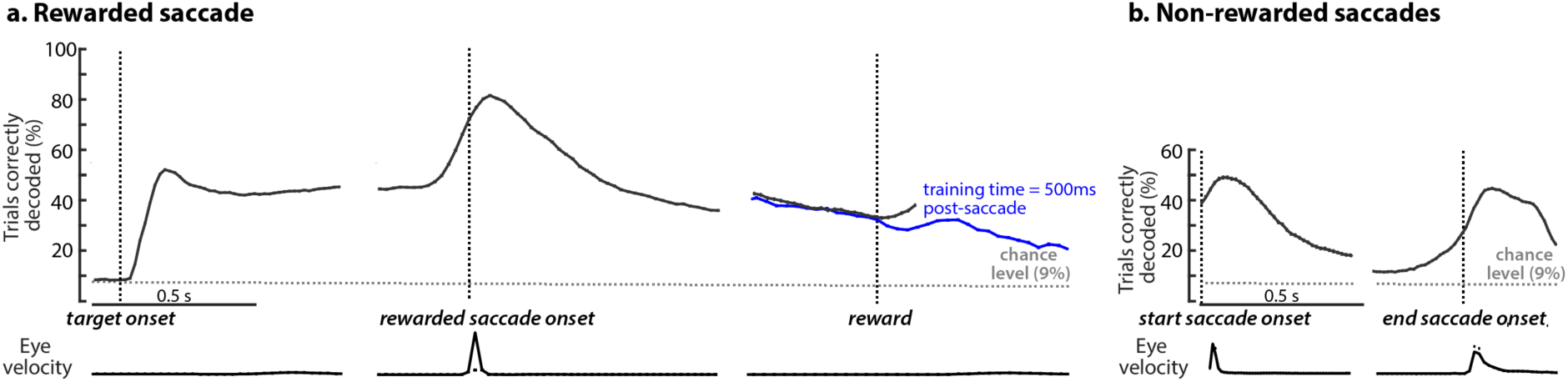
Tuned post-saccadic activity follows every saccade. **a.** Time-specific decoding of the direction of the rewarded saccade at times aligned to target onset, saccade onset and reward. At each time (horizontal axis), a separate multi-class decoder (Linear Discriminant decoder) is trained to predict the direction of the rewarded saccade based on the population response. The vertical axis shows 10-fold cross-validated decoding performance. Decoders are trained and tested on 11 classes (all directions of rewarded-saccades during one session) meaning that chance performance is 9%. The lower panel displays the averaged eye velocity, showing stable fixation prior and post-saccade execution. The blue line indicates decoding accuracy of a single decoder, trained at 500ms post-saccade, and evaluated at many times post-reward. Post-reward there are no constraints on the monkeys’ eye-movements, so here we only show trials where monkeys happen to fixate for longer intervals prior to the next trial. **b.** Cross-validated decoding accuracy when applying the decoders identified for the rewarded saccade (**a**.) to responses aligned to the start (left panel) and end (right panel) saccade. Decoding accuracies are averaged across all sessions and error bars indicate s.e.m. across sessions (n=9).

### Tuned post-saccadic activity follows every saccade (Fig. 2)

We begin our analysis by quantifying the representation of saccade direction in single-trial population responses using cross-validated multi-class decoders^41–46^ (Fig. 2). We decoded the direction of the rewarded saccade from population spike counts at particular times relative to the target onset, the saccade onset and reward delivery.

Cross-validated decoding accuracy varies over time—it rises after the target is presented (Fig. 2a, left), peaks after the end of the saccade, persists throughout the target-fixation, and is still high at and after reward delivery (up to 2s after saccade onset). Decoding accuracy late during *central-fixation* and *target-fixation* is comparable (Fig. 2a, right; Suppl. Fig. 3b, g other monkeys; Suppl. Fig. 2e other decoders). Throughout the *central-fixation* period, the execution of the rewarded-saccade, and the *target-fixation* period, decoding errors almost exclusively reflect read-out directions that are immediately adjacent to the true direction (Suppl. Fig. 2a top row). Decoding accuracy remains high well beyond the time of the saccade, an observation unlikely to be accounted for by transient inputs from motor or sensory areas. Like pre-saccadic activity, post-saccadic activity may thus be a form of persistent, internally generated activity^8,11,39^.

Saccade direction can be robustly read out from the population also after the start and end saccades (Fig. 2b, duration of start saccade = 30+-30ms; duration of end saccade = 140+-80ms; Suppl. Fig. 2a, bottom row, and 2b for a finer comparison; Suppl. Fig. 3c, h other monkeys). Critically, Figure 2b shows the accuracy of decoders that were trained only on activity around the *rewarded saccade*, meaning that the same decoders have high accuracy for all three saccades and implying that the population encoding of saccadic activity is largely preserved across different types of saccades. This finding is consistent with representations of saccades in retinotopic coordinates (see also below, Fig. 3d).

Neural population activity in pre-arcuate cortex thus appears to encode the direction of a saccade from long before it occurs (for the rewarded-saccade) to long after it was completed (for all saccade types). However, the interpretation of post-saccadic activity as a representation of the direction of the immediately preceding saccade is complicated by a feature of the task we analyzed, which is common to many similar tasks. Specifically, the direction of the rewarded saccade is both highly correlated with the direction of the saccade that follows it (the *end saccade,* which often brings gaze back to the fixation point; Fig. 3a, left panel: monkey T; Suppl. Fig 3a, f middle panel: monkeys V and C) and perfectly correlated with the location of the post-saccadic fixation, i.e. the target location. Unless these correlations are accounted for, it remains unclear whether post-saccadic activity is best explained as representing the direction of the previous saccade, the direction of the next saccade, or the location of the post-saccadic fixation.

Interestingly, recordings in monkey C reveal strong post-saccadic activity but little pre-saccadic activity (Suppl. Fig. 3g, h; from a more anterior location in dlPFC than in monkeys T and V), suggesting that pre-and post-saccadic activity amount to fundamentally distinct signals. Below we reach the same conclusion by analyzing datasets tailored to disambiguate between different possible explanations of post-saccadic representations in monkey T and V, for which both pre- and post-saccadic activity occur in the recordings (Fig. 3).

### Post-saccadic activity is not pre-saccadic activity for the next saccade (Fig. 3a-c)

Two observations indicate that post-saccadic activity is unlikely to represent a plan of the next saccade. First, Figure 2b implies that the end saccade (unlike the rewarded saccade, Fig. 2a) is preceded by only very weak predictive activity, which occurs immediately prior to its execution. Second, we studied if predictive activity for the end saccade contributes to the strong post-saccadic activity immediately following the rewarded saccade. To this end, we applied a pre-saccadic decoder (defined 150 to 50ms before the rewarded saccade) to activity *following* the rewarded saccade, and assessed whether the decoder-read out is predictive of the direction of the end saccade.

Notably, we take several steps to ensure that the decoder read-out does not simply reflect the correlations between the directions of the *rewarded* and *end saccades*. For one, we evaluated the accuracy of the read-outs separately for single directions of the *rewarded saccade* (rewarded saccades to three contralateral directions are followed by end saccades in many directions and thus suited to test the decoder; Fig. 3a, dashed rectangle). For another, we created a balanced test set by sampling an equal number of trials from each end-saccade direction (Fig. 3a, bottom-right).

With this unbiased approach, we find that through-out much of the target-fixation period a pre-saccadic decoder (blue vertical line, Fig. 3b) cannot be used to predict the direction of the end saccade (solid lines in Fig. 3b, close to chance; colors correspond to the three rewarded saccade directions in the balanced dataset). During the same period, on the other hand, time-specific decoders (from Fig. 2a) do predict the direction of the preceding rewarded saccade well-above chance (dashed lines in Fig. 3b, and Fig. 2a).

**Figure 3.**
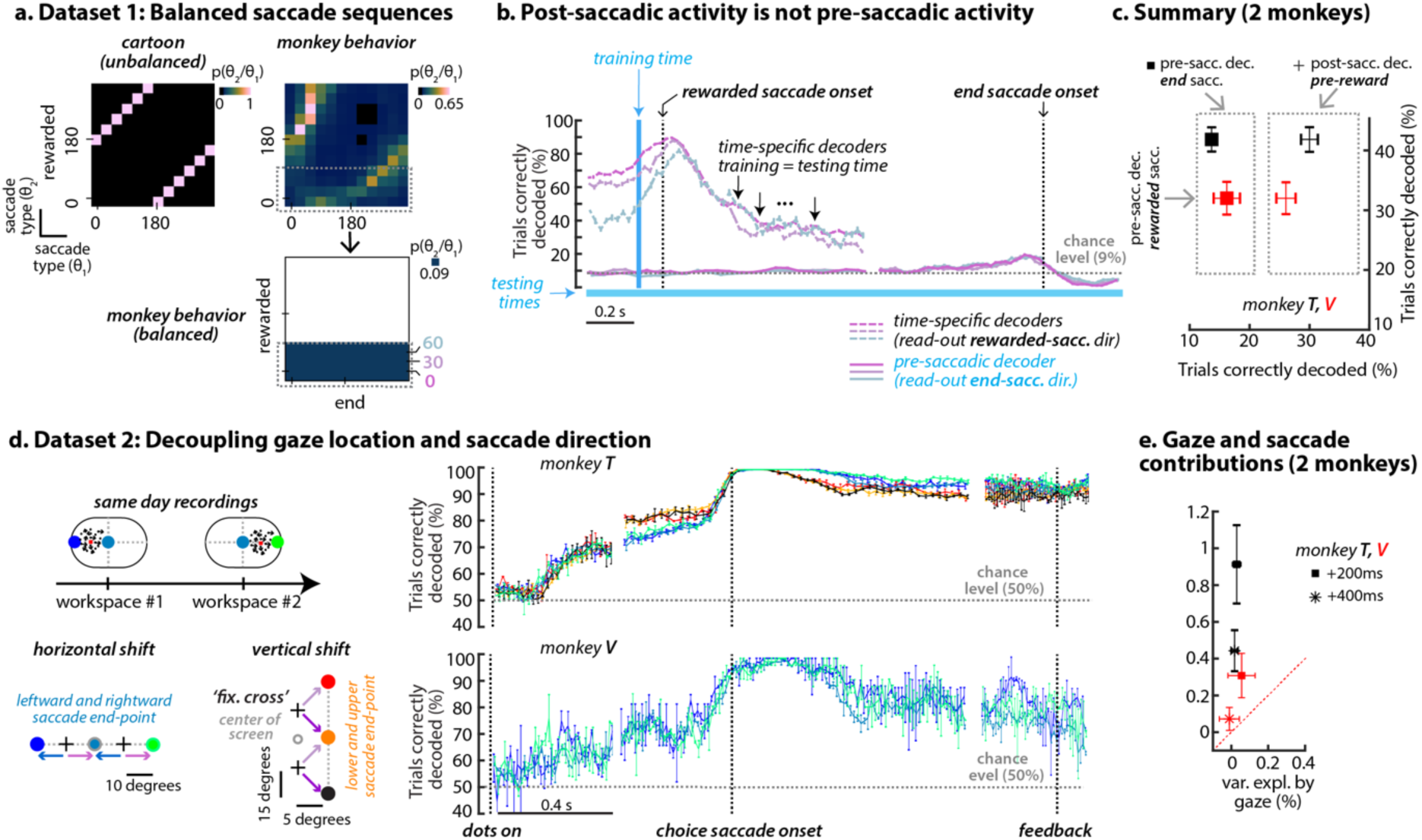
Nature of representations. a-c Modulations of post-saccadic activity due to future actions. Each panel shows histogram of consecutive saccades, expressed as the distribution of directions for the end saccade (columns) conditioned on the direction of the rewarded saccade (rows). For each row, the sum of columns equals 1. Different settings are shown. <u>Left</u> <u>panel</u>: synthetic data for an unbalanced set where all end saccades are directed to the fixation point. <u>Right panel</u>: behavior of monkey T. <u>Bottom panel</u>: a balanced dataset obtained by resampling trials with rewarded saccades towards 0°, 30°, and 60° to obtain a uniform representation of end saccade directions. **b.** We apply a decoder trained on responses prior to rewarded saccade (blue area) to activity after the rewarded saccade separately for the three conditions in the balanced dataset. The resulting read-out is not predictive of the direction of the **end saccade** (bottom three curves). In contrast, on the same set of trials, the direction of the **rewarded saccade** can be decoded with high accuracy when using the time-specific decoders from Fig. 2a (top three curves). **c.** Summary plot. The pre-saccadic activity of the end saccade is much weaker both compared to the post-saccadic activity of the rewarded saccade prior to reward (compare squares and crosses along the horizontal axis) and to the pre-saccadic activity of the rewarded saccade (squares are above the unit line). (n=9) **d-e Non-retinotopic modulations of saccade-related activity. d.** Recordings from a random-dots task that included trials from two “shifted” workspace, whereby the location of the fixation point in a given workspace was shifted either along the horizontal midline (cold colormap) or along the vertical line (warm colormap). For both horizontal and vertical shifts in workspace, SVM decoders of choice-direction achieve comparable, high performance for all gaze locations, including locations corresponding to the end point of saccades with opposite directions (horizontal shift: light-blue curves vs. other cold colors; vertical shift: orange curves vs. other warm colors). Monkey V has sessions only from the horizontal “shifted” workspace. **e.** Summary plot of saccade and gaze modulation at single-unit level. We modeled the activity of each unit with a regression model including linear and non-linear terms for direction and gaze. Saccade direction modulates a larger portion of variance compared to gaze. (n=4, 2 per each shift). Across all panels, error bars indicate s.e.m. across sessions.

We obtained similar results in both monkey T and V (Fig. 3c). Immediately before the onset of the end saccade, predictive activity for the direction of the upcoming end saccade (Fig. 3c, horizontal axis, squares) was weak compared to the representation of the direction of the preceding rewarded saccade (Fig. 3c, horizontal axis, crosses). Together, the above observations imply that post-saccadic activity, consistently in all monkeys, does not represent the plan of a future action.

### Post-saccadic activity does not encode the momentary gaze location (Fig. 3d)

Two observations indicate that momentary gaze location^43–45,47–52^ is also unlikely to be the main contributor to post-saccadic activity. A first indication is given by the finding that post-saccadic decoders trained on the rewarded saccade can also decode the directions of the start and end saccades (Fig. 2b, times following saccade onset). Unlike for the rewarded saccade, for these saccades, the correlation between saccade direction and post-saccadic gaze location is reduced or absent. All start saccades, in particular, end on the central fixation point, meaning that the post-saccadic gaze location is identical across all trials. Yet, the direction of the preceding saccade can be decoded with high performance also following start saccades (Fig. 2b, left).

We find further evidence that post-saccadic does not primarily represent gaze location in a separate behavioral task, for which we partially decoupled the direction of the rewarded saccade and post-saccadic gaze-location. Each experiment included trials from two “shifted” workspaces, whereby the location of the fixation point in a given workspace was shifted either to the left or right from the horizontal midline, or above or below the horizontal midline (Fig. 3d). As a result, one of the choice targets in this task (Fig. 3d, light-blue and orange target) was reached with saccades having very different metrics across workspaces.

Even when post-saccadic gaze location is controlled in this way, the direction of the rewarded saccade can be decoded with high accuracy throughout the central-fixation, movement, and target-fixation periods (right panel in Fig. 3d for monkey T and V). In particular, decoding accuracy is high even on trials that all shared the same post-saccadic gaze location (Fig. 3d, light-blue and orange) and similar to the accuracy on trials where direction and gaze-location covaried (Fig. 3d, other colors). This observation alone implies the existence of a strong representation of saccade direction that is independent of any concurrent representation of gaze location. We further quantified the influence of saccade direction and gaze location at unit-level with a linear regression model, whereby each unit’s activity is captured as a combination of these two factors. Overall, the previous saccade direction explained a substantially larger fraction of the variance in activity than gaze location^29^ (Fig. 3e).

The above findings also make it unlikely that post-saccadic activity represents a gaze-dependent visual input. In fact, selectivity to visual inputs is unrelated to post-saccadic selectivity at the unit-level (Suppl. Fig. 7c). The most parsimonious interpretation of post-saccadic activity is that it represents a retrospective signal, a short-term “memory” of the preceding saccade. The strength and time-course of this action memory appears to vary across saccades, as decoding accuracy differs between different types of saccades (Fig. 2a and 2b; Suppl. Fig. 3b, c and g, h). These differences may imply that post-saccadic activity is modulated by contextual influences, like the temporally discounted reward-expectation^53^ associated with each saccade.

**Figure 4.**
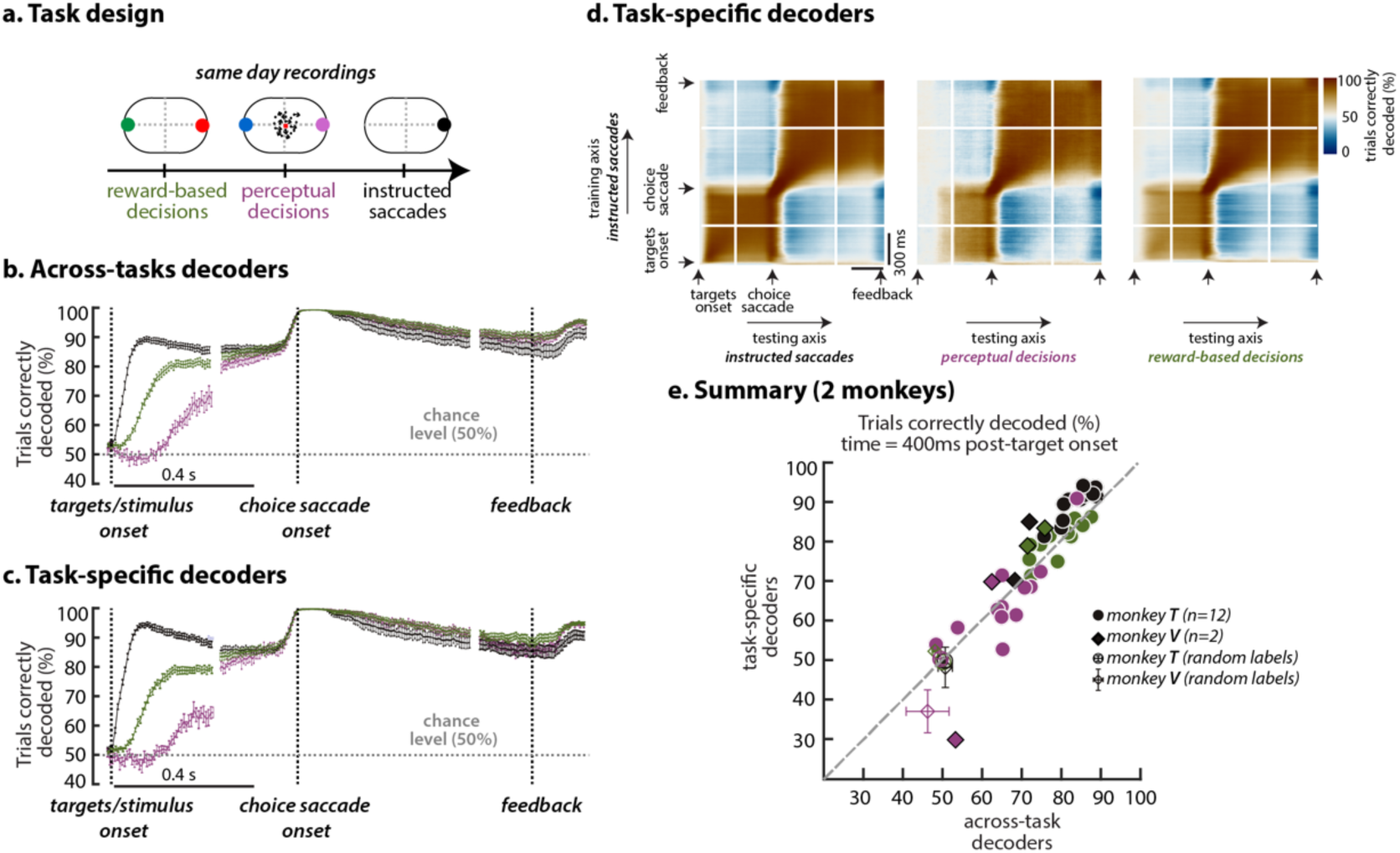
Different tasks, same patterns of activity. Decoding the direction of the rewarded-saccade across different tasks using a binary SVM decoder. **a.** Same-day recordings with common choice targets for reward-based decisions (green), perceptual decisions (purple) and instructed saccades (black). **b.** We decoded choice-direction using a binary SVM on a balanced set across the three tasks. The decoders achieve high accuracy, similarly across all tasks on responses aligned to saccade onset and feedback, but differentiate largely on responses aligned to the appearance of the relevant visual stimulus. Decoders are trained and tested on trials where behavior performance is matched across the three tasks (rewarded trials with high motion coherence in random-dots task and “post-win” trials in the associative task). **c.** We identify decoders for choice-direction for each task (task-specific decoders). Time-course of decoding accuracy is identical to the time-course of the common decoders in b. **d.** We apply task-specific decoders to activity at different times and different tasks. Specifically, we evaluate decoders specific to the instructed saccades from activity from through-out the trial, to perceptual decisions (middle panel) and reward-based decision (right panel). This analysis further reveals that task-specific decoders are, in fact, common across tasks. Specifically, middle and right panel resemble left panel, where training and testing are both on activity from instructed saccades. Decoding accuracies are averaged across sessions and error bars indicate s.e.m. across sessions (n=12). Analogous figure for monkey V (Suppl. Fig. 4). **e.** Summary decoding accuracy for both monkeys at 400ms post target/stimulus onset. Empty markers indicate averaged decoding accuracy when choice labels were shuffled. Decoding accuracies of choice exceed chance levels (50% or empty markers) both for across-task and for task-specific decoders.

### Prospective and retrospective representations have different task-selectivity (Fig. 4)

We further studied context-dependent modulations by comparing the activity of the *rewarded saccade* across three tasks placing different demands on the activity bridging stimuli to actions and rewards (Fig. 4 for monkey T, Suppl. Fig. 4 for monkey V): instructed saccades (as Fig. 1, but with only 2 target locations), perceptual decisions (as in Fig. 3d, but for a single workspace) and reward-based decisions (discussed in detail in the last section of the results). The latter task required monkeys to track which of two colored targets was rewarded at a given time, meaning that choice on a given trial depended on the choice and outcome on the previous trial^24,54–56^.

**Figure 5.**
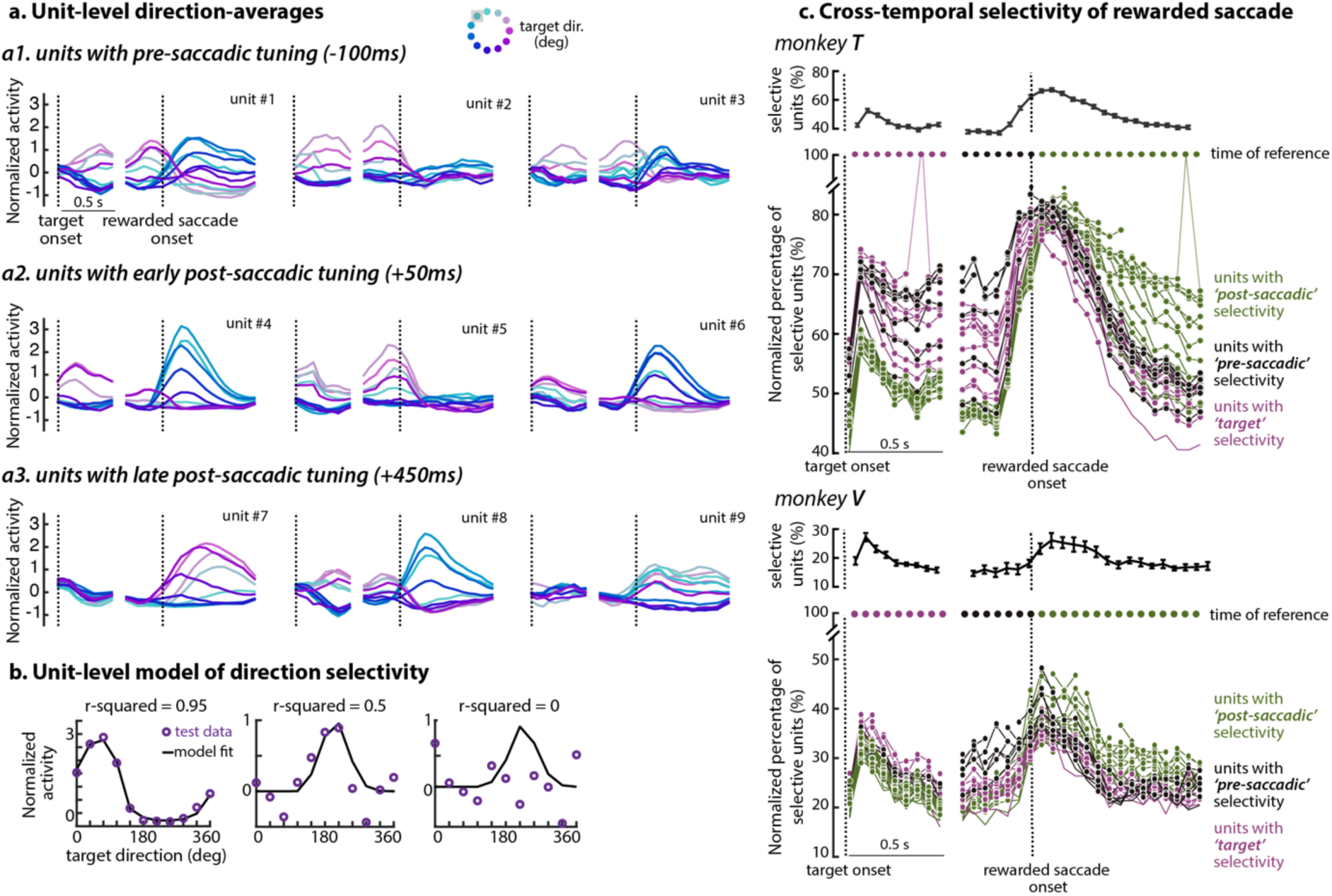
Saccade-related activity in prefrontal units. **a.** Condition-averaged responses of individual units. Example responses for units with high goodness-of-fit (r-squared) **before** (a1), **immediately after** (a2) and **long after** the saccade (a3) are averaged and colored according to the direction of the rewarded-saccade (target eccentricity is ignored). Some units show substantial modulation both before and after the saccade (units #1 and #4). Black dashed line indicates event onset. Grey area in legend of “target dir.” indicates the direction that was not present for this experiment. **b.** Three example fits of the model of direction selectivity to condition-averaged responses with different selectivity levels: from left (highly selective - high r-squared) to right (not selective - r-squared = 0). Model fitting was done through cross-validation. **c.** Time-dependent (top row) and cross-temporal (bottom row) selectivity for the rewarded saccade. Top row: Percentage of selective units (r-squared > 0) on responses aligned to target onset (left) and rewarded saccade onset (right). Bottom row: Colored curves show the percentage of units that are selective both at a reference time (circles on top, color indicates trial epoch - purple, black and green for post-target, pre-saccade and post-saccade, respectively) and at other times in the trial (horizontal axis). For each curve, the lines connecting the corresponding reference time and the two immediately adjacent times (dashed) are mostly omitted. Percentage of selective units is normalized with respect to the percentage of selective units at the reference time, i.e. 70% indicates that 70 percent of the units that are selective at reference time ti are also selective at a different time tj. Circles indicate significant cross-temporal selectivity. Chance level is computed for each time-pair by constructing a null distribution from 1000 permutation tests, where we shuffle the order of units at the two times and compute the overlapping percentage of selective units. Cross-temporal selectivity is considered significant if it exceeds the 95th percentile of this null distribution.

We obtained recordings from all three tasks on the same day, whereby the location of the choice targets was fixed across tasks. We analyzed activity starting from the onset of the visual stimulus that guided the monkeys’ choices, i.e. the target onset in instructed saccades and reward-based decisions, and the onset of the random-dots in the perceptual decisions (Fig. 4b, Suppl. Fig. 4b). To study potential contextual modulations of the underlying representations, we estimated and compared choice-decoders that were *common* across tasks with decoders that were *task-specific*.

Early choice-predictive activity along *common* decoders was strongly modulated by task-context (Fig. 4b, Suppl. Fig. 4b). Predictive activity developed quickly for instructed saccades (black), more slowly for reward-based decisions (green), and slowest for perceptual decisions (purple). We observed these differences even though here we only analyzed trials that were matched for average performance across tasks (e.g. only high-coherency motion trials in the perceptual decisions). In contrast to the strong task-dependency at trial onset, choice-related activity around and following saccade onset was not or only weakly modulated by task-context.

These differences in decoding accuracy at trial onset are observed also when using *task-specific* decoders (Fig. 4c, Suppl. Fig. 4c). The observed task-dependency thus reflects true differences in the strength of the corresponding pre-saccadic representations, as opposed to simply reflecting a “sub-optimal” decoder that captures patterns of activity that are common across tasks, but may not be optimal for some individual tasks. This conclusion is also supported by directly comparing the temporal dynamics of the task-specific decoders^7,17,32,41,57–59^ (Fig. 4d, Suppl. Fig. 4d). Specifically, we applied the decoders from one task, trained at any given time in the trial, to activity recorded at *all times* either in the *same task* (Fig. 4d left) or in a *different task* (Fig. 4d, middle and right). This analysis revealed that early choice related activity is largely explained by a single, stable component that is similar across tasks, but emerges later in the perceptual and reward-based decisions (Fig. 4d middle and right vs. left; compare to Supp. Fig. 4d). Later peri- and post-saccadic activity instead undergoes essentially identical dynamics in all tasks (Fig. 4d). Consistently in both monkeys (Fig. 4e), choice representations thus transition between the same patterns of activity in all tasks, albeit with somewhat different speeds (Fig. 4b).

### Prospective and retrospective signals are mixed in individual units (Fig. 5)

Having established that dlPFC populations maintain prospective and retrospective representations of saccade direction, we asked how these representations are organized at the unit-level. In particular, prospective and retrospective signals could be maintained by separate populations of neurons, or could be mixed within a single population. To address this question, we focus on the instructed-saccade task shown in Fig. 1. An examination of example units shows substantial variability across the population in the temporal dynamics of saccade-modulated activity^8,11,19,36–39,60^ (Fig. 5a), with some units selective prior to saccade (units #2 and #5), after the saccade (units #7 and #9) or both before and after the saccade (units #1 and #4).

To quantify the strength and dynamics of directional selectivity in individual units, at any given time in the trial we fitted a bell-shaped function to the activity averaged by target direction^11^, while ignoring target eccentricity (Fig. 5b). We considered a unit to be direction selective at a particular time if the cross-validated r-squared value of the corresponding fit was higher than 0, i.e. the model describes the direction-averaged responses better than a constant. For selective units, we then defined the *preferred direction* as the corresponding model parameter (the peak location of the fitted tuning curve).

**Figure 6.**
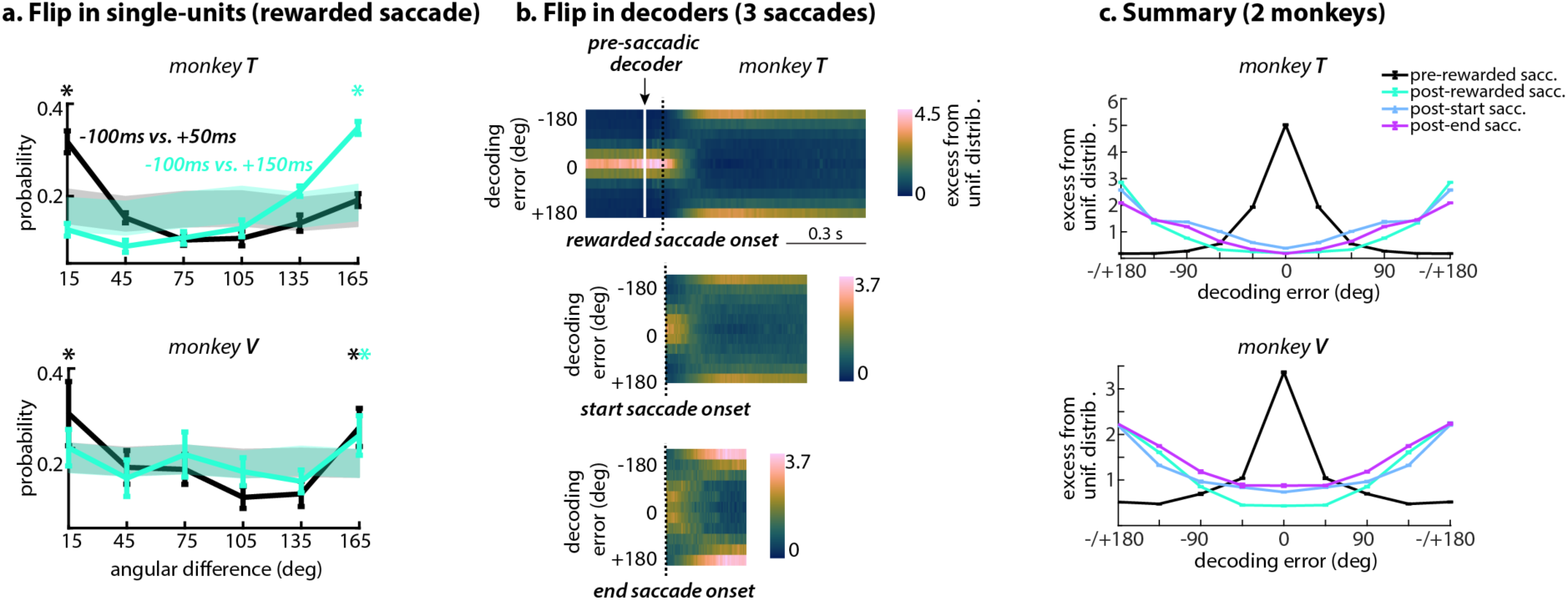
The relation between pre and post-saccadic activity. **a.** Selectivity dynamics of units with cross-temporal tuning. Plot shows a histogram of angular difference between the pre-saccadic and post-saccadic preferred directions units that are selective at both times. We construct a null-distribution by shuffling the order of units and thus assuming no relation between a unit’s preferred direction at pre and post-saccadic times. The shaded area marks the 5^th^ and 95^th^ percentile of this null-distribution over 1000 random repetitions. The distribution of angular differences is binned in 30 degrees bins. At +50ms, the post-saccadic and pre-saccadic preferred directions tend to match (peak of distribution at 15° - [0: 30] degrees, “stable”), but at +150ms the preferred directions have mostly flipped (peak of distribution at 165° - [150: 180] degrees, “flip”). **b**. Time-dependent histograms of decoding errors of a pre-saccadic decoder (trained on responses from -150ms to -50ms prior to rewarded saccade, Linear Discriminant decoder) when applied onto post-saccadic responses of the rewarded saccade (top), start saccade (middle) and end saccade (bottom). Each histogram (vertical axis) is normalized with a uniform distribution, where the uniform distribution is estimated by computing the angular difference between two random draws of N angular discrete values matching the target locations, 1000 times. The y-axis shows the empirical distribution divided by the mean of the estimated uniform distribution. Dashed black line indicates saccade onset. **c**. Summary plot illustrating the distribution of decoding errors of a pre-saccadic decoder applied to pre-saccadic responses (black line, responses from -150ms to -50ms prior to rewarded saccade) and post-saccadic responses (colored lines, responses from 250ms to 350ms post rewarded, start and end saccades). Decoding errors are shown only for saccades directed towards the contralateral hemifield, because of their strong pre-saccadic activity (Suppl. Fig. 2d for monkey T, Suppl. Fig 3b for monkey V). For each monkey, we use the decoder with the highest decoding accuracy for the direction of rewarded saccade (See Suppl. Fig. 2e for monkey T and Suppl. Fig. 3e for monkey V), but results were quantitively similar across other decoders. All figures contain results averaged over sessions. (n = 9 for monkey T and n = 10 for monkey V). Error bars indicate s.e.m. across sessions.

We find that a substantial fraction of units encodes direction at any given time in the trial in all monkeys (Fig. 5c, top for monkey T and V, Supp. Fig. 7b for monkey C). The fraction of selective units varies throughout the trial, largely mimicking the time-course of the population-level decoders (Fig. 2; Supp. Fig. 3b, g for monkey V and C). To compare the strength of tuning in individual units across time, we defined a “cross-temporal selectivity” measure (Fig. 5c, bottom row), which quantifies the percentage of units that are direction selective at a given reference time (small circles on top and curve of the corresponding color) as well as at a different comparison time (horizontal axis).

The cross-temporal selectivity is broadly consistent with mixed selectivity^59^. A substantial fraction of units that are selective after the saccade are also selective before the saccade or right after target onset (about 50% and 25% in monkeys T and V, Fig. 5c, green curves; circles indicate significant cross-temporal selectivity). Similarly, many units that are selective after the target onset or before the rewarded saccade are selective also long after the saccade (Fig. 5c, purple curves).

### Signal mixing within units is not random (Fig. 6)

The relation between prospective and retrospective representations in individual units is highly structured. In units showing both pre- and post-saccadic tuning (Supp. Fig. 7a), we computed the angular difference between the preferred direction estimated immediately before saccade onset (-100ms) and two times following the end of the saccade (+50ms and +150 after saccade onset). In monkeys T and V, more units than expected by chance show an angular difference close to 180 deg, implying that the preferred direction tends to “flip” between pre- and post-saccadic activity (Fig. 6a).

The flip in preferred direction is also prominently reflected in the inferred population decoders. Applying a pre-saccadic decoder trained during the central-fixation to the activity in the post-saccadic epoch results in a pattern of read-out errors strongly biased towards the direction *opposite* to the true saccade direction (Fig. 6b, upper row, pre-saccadic decoder; decoding error = 180°). The bias is strongest shortly after completion of the rewarded saccade, but persists throughout even the longest target-fixations (Suppl. Fig. 2c).

Similar read-out errors are observed when applying the same pre-saccadic decoder to the post-saccadic activity of both the start saccade and end saccade (Fig. 6b, middle and bottom rows). Crucially, the prominent regularities in the metrics of saccades that follow the rewarded saccade (Suppl. Fig. 6 for monkey T; end saccades tend to be opposite to the rewarded saccade; Suppl. Fig. 3a for monkey V) are largely absent for the start and end saccades (Suppl. Fig. 6, left and right) implying that the inferred structure between pre and post-saccadic selectivity is not simply a consequence of these regularities in the behavior.

Overall, the relation of pre-saccadic and post-saccadic responses is far from random (Fig. 6c, summary for monkey T and V), but rather reveals a highly structured way for the neural population to transition from representing the plan of an action to representing its memory. These structured representations stand in contrast to the findings of prior studies showing that many abstract variables are randomly mixed across units^7,17^, implying potentially different encoding strategies for spatial and abstract variables.

### An associative learning task (Fig. 7)

We studied how the identified neural representations change though-out learning in an associative learning task^24^, a kind of task that was previously shown to rely on an intact pre-arcuate gyrus^61^. Monkeys were engaged in a two-alternative, forced-choice task that required them to track which of two targets (red or green) was being rewarded at any given time (Fig. 7a). Because the timing of switches in rewarded color was unpredictable, the optimal strategy is “win-stay, lose-switch”^54–56:^ if a given color was rewarded (“win”), the monkey should choose the same color again on the next trial (“stay”). Instead, after a choice that was not rewarded (“lose”) the monkey should switch to the other color (“switch”). Monkeys’ performance gradually improved over the course of many weeks of exposure to this task (Fig. 7b).

Achieving optimal performance in this task requires both fast and slow learning^62^. On the fast time-scale of trials, monkeys must update their beliefs about what color and location will be rewarded on the current trial based on the actions and outcomes on the preceding trials. On the slow times-scales of days and weeks, monkeys must infer the task rules and learn a strategy to optimally harvest rewards. We characterized fast learning with logistic regression models fit to the behavior in a single session, and then studied how the corresponding strategies are shaped by slow learning across sessions (Fig. 7c). We separately modeled the influence of the (task relevant) target color and the (task irrelevant) target location on the monkeys’ choices (Fig. 7c, circles and squares) and their interaction with previous outcome (win or lose, x- and y-axes).

**Figure 7.**
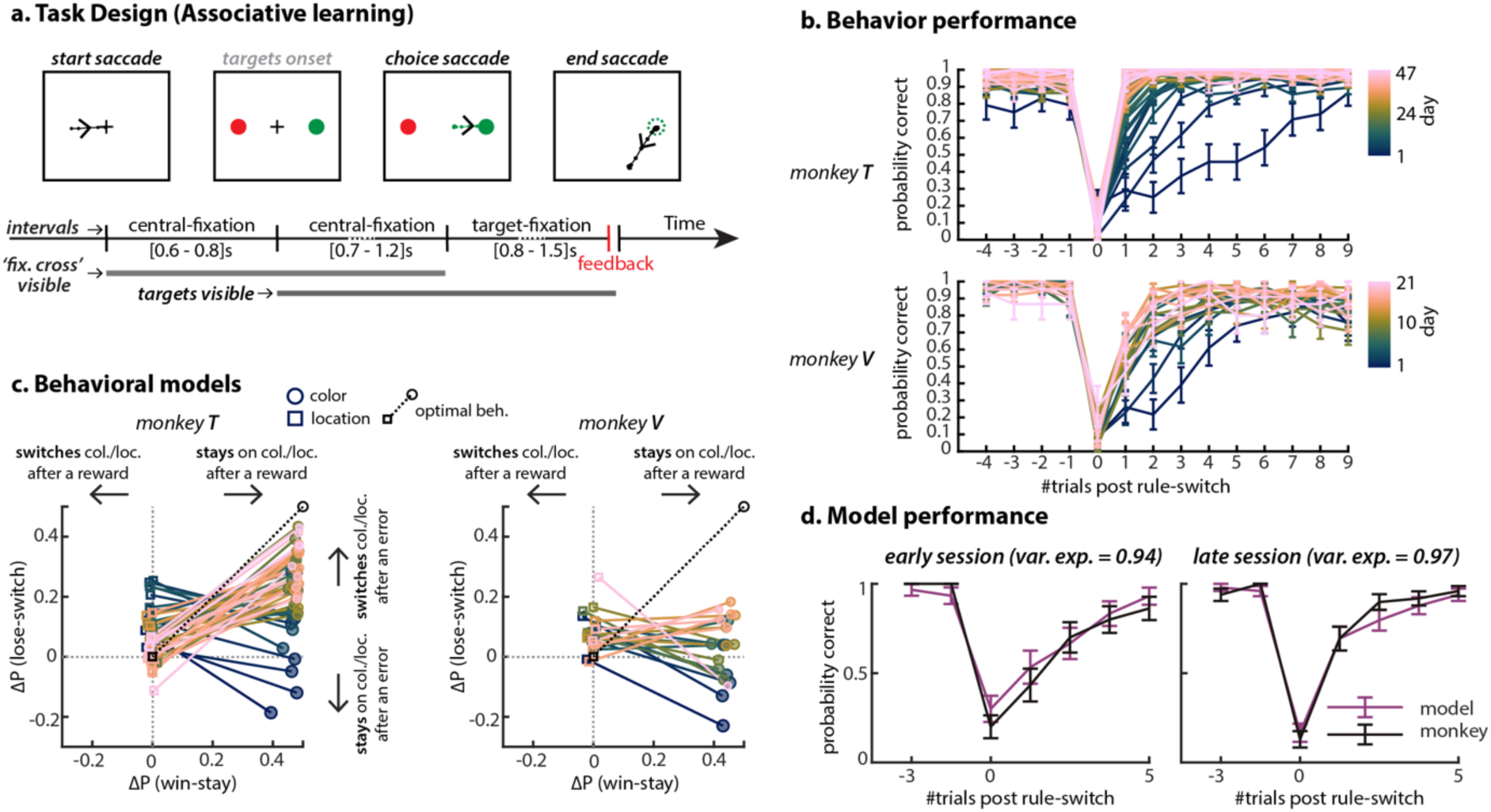
An associative task and behavior performance. **a.** Subjects performed an associative task that had the same trial structure as the instructed saccade task, namely a central-fixation period of random duration between targets appearance, and the saccade (choice-saccade) towards one of them; followed by a target-fixation period of random duration requiring monkeys to fixate the chosen target until feedback. On any given trial, the reward contingencies depend on the outcome and the chosen color of the previous trial. The mapping between color and location is random from trial-to-trial. **b.**Behavior performance varies substantially on trials following a rule-switch. This implies that the monkeys’ behavior after errors changes through-out learning. In contrast, the behavior after rewards is almost constant. **c.** Behavioral models describing the strategies the monkeys use to harvest rewards. ‘Win-stay - lose-switch’ behavior is modeled with logistic regression using either the relevant history (choice-color) or the irrelevant history (choice-location). Figure shows the estimated probability that the monkeys will either choose the same choice-color/choice-location as in the previous trial following rewarded trials (win-stay, horizontal axis) or switch following error trials (lose-switch, vertical axis). From the estimated probabilities we subtract the simulated probabilities of a random strategy. Thus, for an optimal agent (black cartoon) ΔP for location is 0 and for color is 0.5. **d.** Simulated color-choices using the model of previous relevant history explain well behavior around rule-switches, both early and late in training.

In both monkeys, slow learning primarily involved changes in how monkeys reacted to unrewarded trials (Fig. 7c, y-axis). Monkeys learned to consistently switch colors after a lose trial (Fig. 7c, circles; ΔP(lose-switch) gradually approaches 0.5). Notably, monkeys consistently stayed on the rewarded color already during the very first session on this task (Fig. 7c, ΔP(win-stay) close to 0.5). Likewise, monkeys’ choices were more strongly affected by the irrelevant target location after lose trials compared to win trials (Fig. 7c, squares; larger differences from 0 along y-compared to x-axis). Notably, in monkey T learning also involved overcoming an initial (incorrect) spatial strategy (Fig. 7c left, squares: ΔP(lose-switch) gradually approaches 0.5). In both monkeys, the inferred stay and switch probabilities were sufficient to explain the dynamics of performance after a change in the rewarded color, implying that fast learning primarily relies on information from the immediately preceding trial.

**Figure 8.**
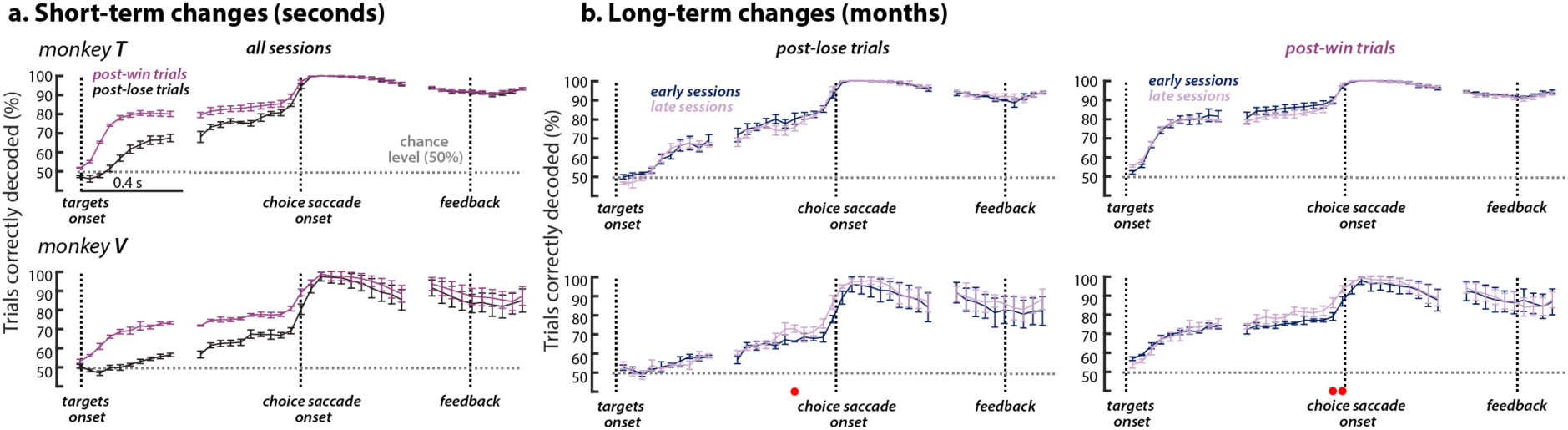
Choice-representations during learning. **a.** Decoding accuracy of saccade direction through-out the trial. Decoders are trained only on post-win trials, i.e. trials following rewarded trials, and evaluated on (1) post-lose trials, i.e. trials following unrewarded trials, and (2) on a held-out set of post-win trials. We resample trials to have the same train and test set size across training days. Choice-related activity is substantially modulated by the outcome of the previous trial at targets onset, but not at times relative to saccade onset. Decoding accuracy is averaged over all sessions and error bars indicate s.e.m. (n=36 for monkey T and n=21 for monkey V). **b.** Decoding accuracy of post-lose trials split for early and late training days. Early sessions are considered the first half and late sessions the second half. Dots on the bottom indicate p-value < 0.05 as estimated from a partial correlation that correlates training half (1 for early and 2 for late) and decoding accuracy of each session. We use partial correlation to control for any effects that radius (continuous values) and target configuration (one-hot encoding) might have on decoding accuracy. **c.** Same as b. but for post-win trials.

### Neural changes across fast and slow learning (Fig. 8)

We studied how choice representations relate to both fast and slow learning by comparing decoding accuracy for the direction of the choice saccade (choice location) across trials differing in the outcome of the immediately preceding trials (Fig. 8a; fast learning) and across early and late sessions (Fig. 8b; slow learning). Despite its task relevance, choice color was only weakly represented in the recorded areas (Suppl. Fig. 9).

The recorded choice-representations were modulated by trial-history, albeit only in pre-saccadic activity. Post-lose trials had weaker choice predictive activity compared to post-win trials up until saccade execution. After the saccade, choice-related activity instead was equally strong across both trial types. The observed history-dependence of pre-saccadic activity seems consistent with the above finding that slow learning primarily involves adjustments to the strategy following lose trials (Fig. 7c) and may reflect low confidence, frequent changes of mind^63^, or a slowing down of neural responses after negative feedback^64,65^.

Notably, the monkeys’ overall task performance did not correlate with any of the features of the responses we considered (Fig. 8b). Decoding accuracy on post-lose or post-win trials was similar in early and late sessions during all trial epochs, despite the monkeys performing closer to optimal on late vs early sessions (Fig. 7c). The representation of other task variables, like choice color, was also unchanged throughout learning (Supp. Fig. 9b). Overall, the properties of pre- and post-saccadic representations thus appeared to be largely fixed over long time-scales.

## Discussion

We measured and quantified population activity in pre-arcuate cortex of macaque monkeys across different tasks (instructed, perceptual discrimination and reward-based decisions), different type of saccades (instructed and free) and different learning time-scales (seconds and months) to obtain a comprehensive characterization of saccade-related activity in pre-arcuate cortex in different contexts.

### Properties of pre- and post-saccadic activity

We find that post-saccadic activity is the strongest and most consistent form of saccade-related activity, both at unit-level (Fig. 5c, top row for monkey T and V, Suppl. Fig. 7b for monkey C) and at population-level (Fig. 2 for monkey T, Suppl. Fig. 3b, g for monkey V and C). The direction of the rewarded saccade can be best decoded after the saccade is already completed, and decoding performance remains high until the time of feedback, throughout a delay period during which the gaze is fixed (Fig. 2a). This finding is notable in an area of dlPFC that was previously primarily associated with pre-saccadic responses^66^. A key factor in revealing the prominence of post-saccadic activity was to more closely balance the duration of pre- and post-saccadic epochs in our tasks.

The persistence of post-saccadic activity seems at odds with the findings of some previous studies, which instead reported largely transient activity^8,30^. These past studies, however, did not include a temporal separation between the saccade and the feedback. The persistent nature of post-saccadic activity might become apparent only when such a delay period is included in the task, since any task-relevant saccade-related information may have to be maintained until feedback is provided.

As in more posterior areas of PFC^8,11,18,19^, but not more anterior ones^27,28^, post-saccadic activity is intermingled with pre-saccadic and movement related activity (see Suppl. Discussion on retinotopic coordinates). When pre- and post-saccadic activity co-occurs in single neurons, they are tightly linked—the preferred direction typically flips by 180 degrees between the pre- and post-saccadic epochs for all saccades (Fig. 6). Analogous flips in selectivity have been observed before in relation to saccades^8,11^, but may also occur in other settings^32,67,68^. This structure stands in contrast with the common finding that in associative areas task-related variables are often randomly mixed in the population^7,17,59^. As discussed below, one possible function of the observed flips may be to update a representation of visual space across saccades.

Pre- and post-saccadic activity are differently modulated by saccade type and task. Unlike pre-saccadic activity^11,18,38,69^ (but see^70^), post-saccadic activity occurs after every saccade, but is strongest and lasts the longest following “rewarded” saccades (i.e. the last saccades preceding feedback and reward delivery, Fig. 2a). Weaker and more short-lived post-saccadic activity follows the start saccades that initiate a trial, and the end-saccades that follow the reward (Fig. 2b).

During learning, we find that choice representations reflect the recent trial-to-trial history, but not day-by-day behavioral improvements. This was particularly surprising for trials following errors, where the behavior strongly changes through-out the learning process. The fast adjustment to rule-switches in later sessions (pink lines in Fig. 8b) indicate that monkeys correctly interpret errors as informative factors relevant for future decisions. Nevertheless, the weak choice-representations on post-error trials are observed in late and early sessions, despite an almost impeccable behavioral performance during late sessions.

The fact that a linear decoder was capable of reading the monkey’s choice independent of the task (Fig. 4b) suggests that dlPFC constitutes an advanced processing stage where representations are rather rigid and resemble “domain-general”^71^ and/or “untangled”^72^ representations. These representations did not reflect the large changes in behavior occurring while monkeys learned the reward-based decisions. This task learning^73^ could instead reflect plasticity in areas upstream of dlPFC or in recurrent circuits involving both cortical and subcortical areas^74^.

### Possible functions of post-saccadic activity

The finding that post-saccadic activity amounts to an action memory is consistent with a key role of dlPFC in retrospective monitoring of behavioral context and in binding the past to the present^20–22,24,27–29,75,76^ (see Suppl. Discussion for previously proposed functions). The persistent representation of an action memory may be similar to the maintenance of other behaviorally relevant variables in working memory (e.g.^3,7,77^). Representations of past stimuli in dlPFC, however, are not limited to persistent activity, but can remain present in “activity-silent” traces that reappear as activity on future trials^78^. Temporary records of previous actions are required by many reinforcement learning algorithms to evaluate the actions’ relevance with respect to rewards^28,74,79–82^ (see Suppl. Discussion on choice memories^83–86^ and eligibility traces). Neither pre-nor post-saccadic activity was modulated by slow learning processes, suggesting that the underlying spatial representations are a rigid, core feature of dlPFC. Such rigid representations may coexist with the flexible emergence of representations of abstract task variables^7,59,87–89^. The separation of rigid and flexible representations could be computationally advantageous, as it might reduce task-interference^90^ or catastrophic forgetting^90^.

Beyond learning, post-saccadic activity could contribute to updating representations of visual space in PFC and to maintaining visual stability across saccades^19,92,93^. Our own work and previous studies implies that visual stimuli, salient locations, action-plans, and their memories are all represented in dlPFC in maps organized in retinotopic coordinates^11,19,38^. Any behavior requiring more than a single saccade, like the visual exploration of a scene, or the execution of sequences of saccades to multiple remembered locations, requires updating these retinotopic maps following each saccade, a process often referred to as remapping^26,94–98^. Concretely, after a saccade, a retinotopic map needs to be updated by shifting it along a vector that is the exact *opposite* of the vector of the saccade that was just executed (Suppl. Fig. 10 and Fig. 13 in ^19^). The prominent flip in direction selectivity observed after each saccade could quickly^31^ provide such an update signal, or could reflect the outcome of the update process.

A contribution of post-saccadic activity to updating spatial representations would imply a critical role for PFC in predicting and compensating for the consequences of one’s own actions^92,93,99,100^. Consistent with such a role, impairments in generating and incorporating predictions are thought to be a defining feature of schizophrenia^101,102^, which consistently involves prominent changes in prefrontal circuits^102,103^ as well as an impaired ability to generate long and frequent saccades in visual exploration^104^.

## Conclusion

Pre-arcuate cortex actively maintains accurate, persistent representations of saccades before, during, and after each saccadic movement. The representations of saccadic action plans and saccadic action memories are expressed in the same frame of reference, retinotopic coordinates, making them well-suited as a basis for reinforcement learning algorithms^105^ and for the computations underlying visual stability across saccades^92^. The observed, concurrent representations of saccadic action preparation, action execution, and action memories support a prominent role of PFC in linking events across time. An important question for future studies is how the strong and rigid spatial representations we described relate to the flexible representations of more abstract behavioral variables in PFC and throughout the brain^27,28,106,107^.

## Authors Contribution

I.C. and V.M. designed the study and the methods. J.R. conceived and conducted the experiments and collected the data. I.C. performed the analyses, with input from V.M. and assistance from S.K. S.K. provided software for data visualization and data pre-processing. I.C., V.M. and S.K. wrote the manuscript. All authors were involved in discussing the results and the manuscript.

## Acknowledgments

We thank W. T. Newsome for providing us the data and for discussions on the results. We also thank members of the Mante Lab for valuable feedback throughout the project.

## Supplementary discussion

### Retinotopic coordinates

Both pre-saccadic^11^ and post-saccadic representations reflect retinotopic coordinates. Like other types of movements, saccades could in principle be represented in a variety of alternative coordinate systems, from head-centered, to body-centered, and world-centered coordinates^108,109^. Like past studies in dorso-lateral prefrontal cortex^11,110^, we, however, find weak representations of saccades in coordinate systems other than retinotopic coordinates (Fig. 3d, e). Potential candidate structures for such representations in other coordinate systems involve hippocampal areas and parts of PFC closely linked to it^52,111–114^.

### Function of post-saccadic activity

The properties of post-saccadic activity in pre-arcuate cortex appear inconsistent with a number of proposed hypotheses about its function. Since post-saccadic activity is not predictive of the next saccade, it is unlikely to represent a plan for a future action (see also^11^). A hypothesized role in “resetting” activity in PFC, to set the stage for a new saccade plan^30^, seems at odds with the observation that post-saccadic activity can persist over long temporal intervals. This persistence, together with a pronounced dependency on reward expectation, also rules out the possibility that post-saccadic activity represents a corollary discharge for saccades. Instead, post-saccadic activity might be involved in retrospective monitoring the behavioral context^20–22,24,27–29,75,76^.

### Relation to choice sequences

Choice memories represented as sequences of activity in rodent prefrontal and parietal cortex^83–86^ have been proposed as neural correlates of eligibility traces. The post-saccadic activity we report here, however, differs in some respects from previously reported choice related sequences^83–86^. First, we show that post-saccadic activity appears to follow every saccade, not just saccadic movements related to a choice between learned alternatives. Second, post-saccadic activity changes smoothly as saccades direction is varied along a circle (Suppl. Fig. 2a) and can be traced back to single-unit response fields. Third, we observe post-saccadic activity during fixation periods in which task-relevant movements are suppressed, effectively excluding possible explanations of this activity through movement confounds^115^.

### Eligibility traces

Many reinforcement learning algorithms^105^ use eligibility traces, i.e. temporary records of previous actions, to evaluate the actions’ relevance with respect to rewards^28,74,79–82^. Eligibility traces of eye movements are particularly important for learning, as eye movements provide a fast feedback of motor performance^116,117^. Implementing such algorithms in neural circuits is challenging, as learning may rely on biophysical mechanisms like spike-timing dependent plasticity (STDP) that operate on much shorter times-scales than the task-events relevant for behavior^118,119^. Past proposals on how to link synaptic plasticity to times-scale of behavior include tagging synapses to make them eligible for future reinforcement-driven changes^120,121^ or prolonging the temporal footprint of STDP^122,123^. Such mechanisms seem ill-suited to the tasks studied here, as the duration of the target-fixation period separating action and outcome outlasts even the longest-documented windows of adult-brain STDP^124^. On the other hand, by actively representing persistent components originating from different times in the trial in the network, representations of stimuli, actions and outcomes that are separated in time might be made to temporally overlap in the brain, thus allowing learning to occur through fast mechanisms like STDP. Our finding that post-saccadic activity is modulated by *saccade type* may imply that action memories, similarly to working memory of sensory information, are maintained in PFC flexibly, and preferentially for those actions that are most relevant for learning. Such action memories may complement or interact with alternative mechanisms that could allow task-relevant signals to be maintained without persistent activity^78^, such as an “activity-silent” memory emerging from changes in synaptic efficacy^125,126^.

## Supplementary Figures

**Supplementary Figure 1.**
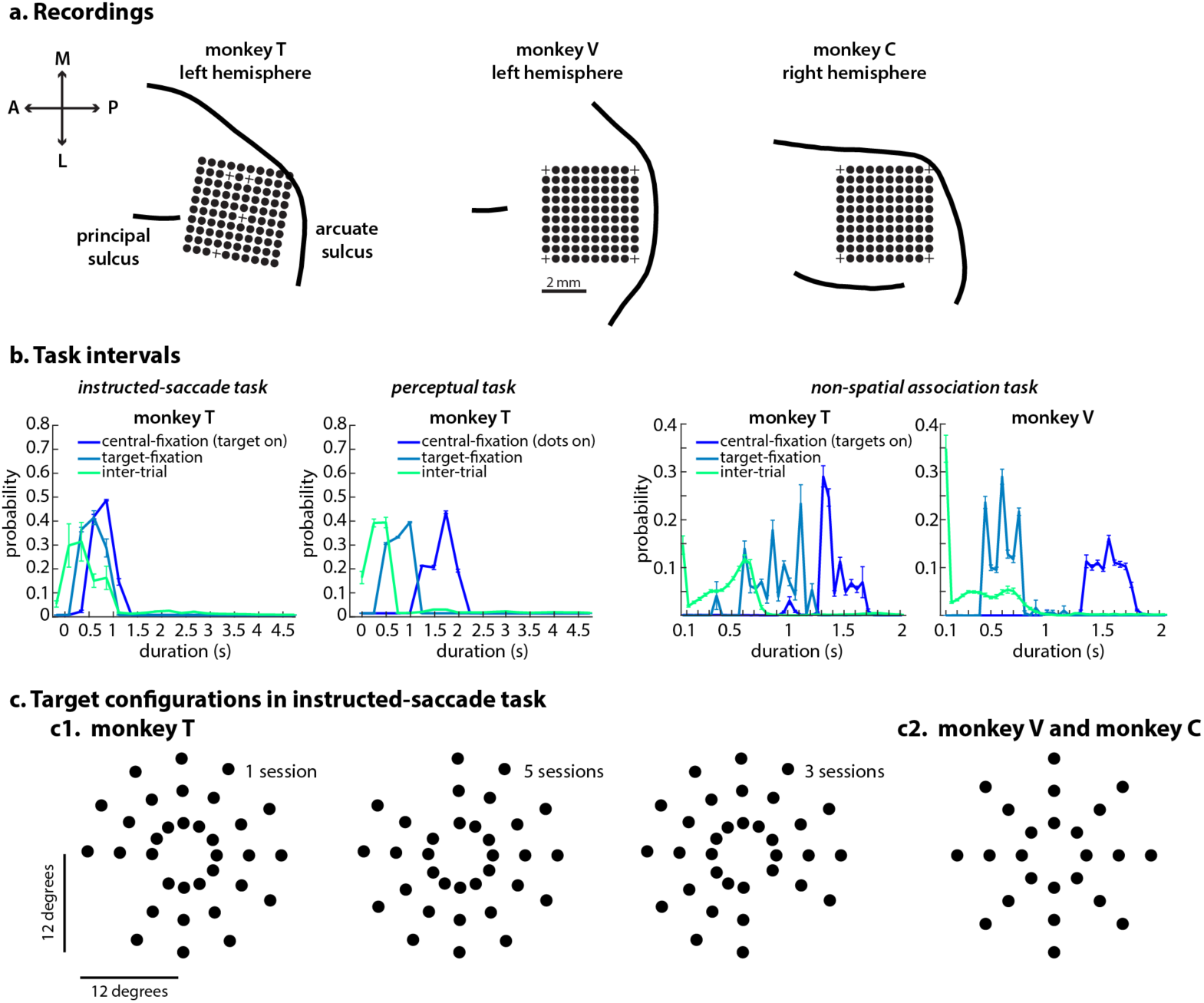
Neural recordings and task parameters. **a.** In all three monkeys, we obtained single-unit and multi-unit recordings from a 10x10 array implanted in pre-arcuate cortex. Black circles indicate the cortical locations of the 96 electrodes used for recordings. **b.** Durations of three intervals occurring in each trial. Central-fixation (target/dots/targets on for instructed/motion-discrimination/reversal learning task): from the onset of the relevant stimulus until the offset of the fixation point. Target-fixation: from the onset of the rewarded saccade until the reward delivery. Inter-trial: from the reward delivery of a given trial until the onset of the fixation point on the next trial. For reversal learning task, intervals for all three monkeys are shown. Note that the length of target-fixation interval is longer for monkey T than for monkey V and C. **c.** Target configurations for the instructed-saccade task - 33 unique locations for monkey T and 24 unique locations for monkey V and C. For monkey T, the angular difference between two neighboring targets is 30 degrees, but on each session, one direction is missing (left panel - 210 degrees, middle panel - 120 degrees, right panel - 300 degrees). For monkey V and C, the angular difference between two neighboring is 45 degrees. For all monkeys, the radial difference between two neighboring targets is 4 degrees. Radii values are 4, 8 and 12 degrees. Error bars indicate s.e.m. across sessions.

**Supplementary Figure 2.**
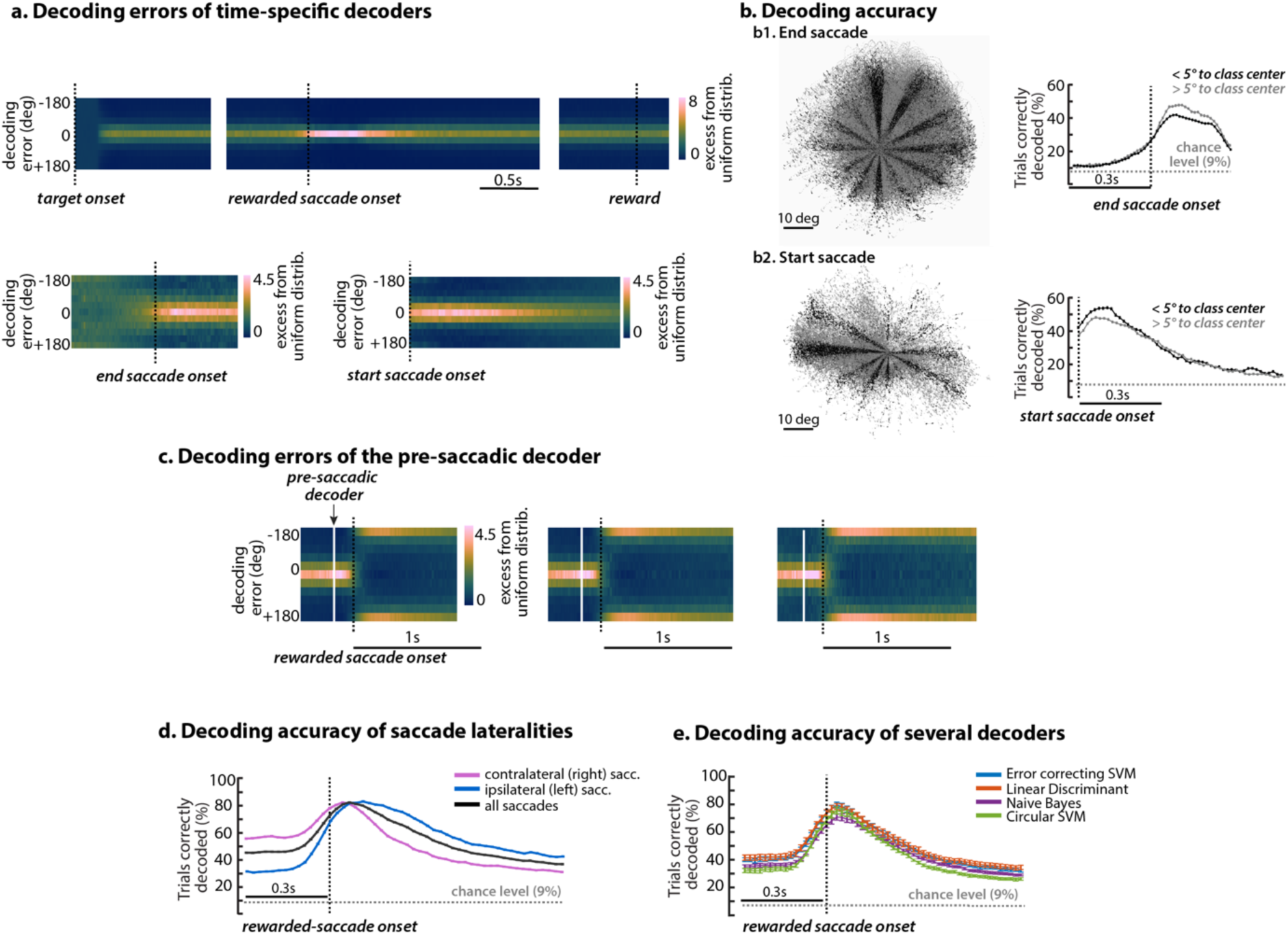
High-dimensional decoders. **a.** Time-dependence of decoding errors; the distribution of angular errors (vertical axis) when testing times (horizontal axis) are chosen to coincide with training times for times relative to target onset (top row, left panel), to rewarded saccade onset (top row, middle panel), reward delivery (top row, right panel), end saccade onset (bottom row, left panel) and start saccade onset (bottom row, right panel). **b.** Eye-movement trajectories (left) and time-specific decoding (right) for end (top) and start saccades (bottom). Unlike for the rewarded saccades, the direction of the end and start saccades is continuous. To apply the decoders trained on the rewarded saccade, we discretize this continuous variable into bins whose centers match the directions of the rewarded saccade. To study how this binning affects the decoding performance, we assigned saccades into two groups: saccades with directions close to the respective category center (left panels, black) and saccades with directions far from this center (left panels, gray). Decoding performance for both groups is similar to performance when all trials are included Fig. 2b. The lower decoding performance of end and start saccades compared to rewarded saccades thus does not seem to be a consequence of the different distribution of saccade directions. **c.** We applied a pre-saccadic decoder to activity before and after the rewarded saccade (as in Fig. 6b, top row). We separated trials based on the duration of the target-fixation period (0.8s, 1s and 1.2s). The angular errors close to 180 degrees reflect a flip in read-out along the pre-saccadic decoder at times following the rewarded saccade. **d**. Decoding accuracy split by saccade laterality. Contralateral saccade have higher pre-saccadic activity than post-saccadic activity, while ipsilateral saccades have higher post-saccadic activity than pre-saccadic activity. **e.** Decoding accuracy of different decoders. Results are averaged over sessions and error bars indicate s.e.m. across sessions (n=9)

**Supplementary Figure 3.**
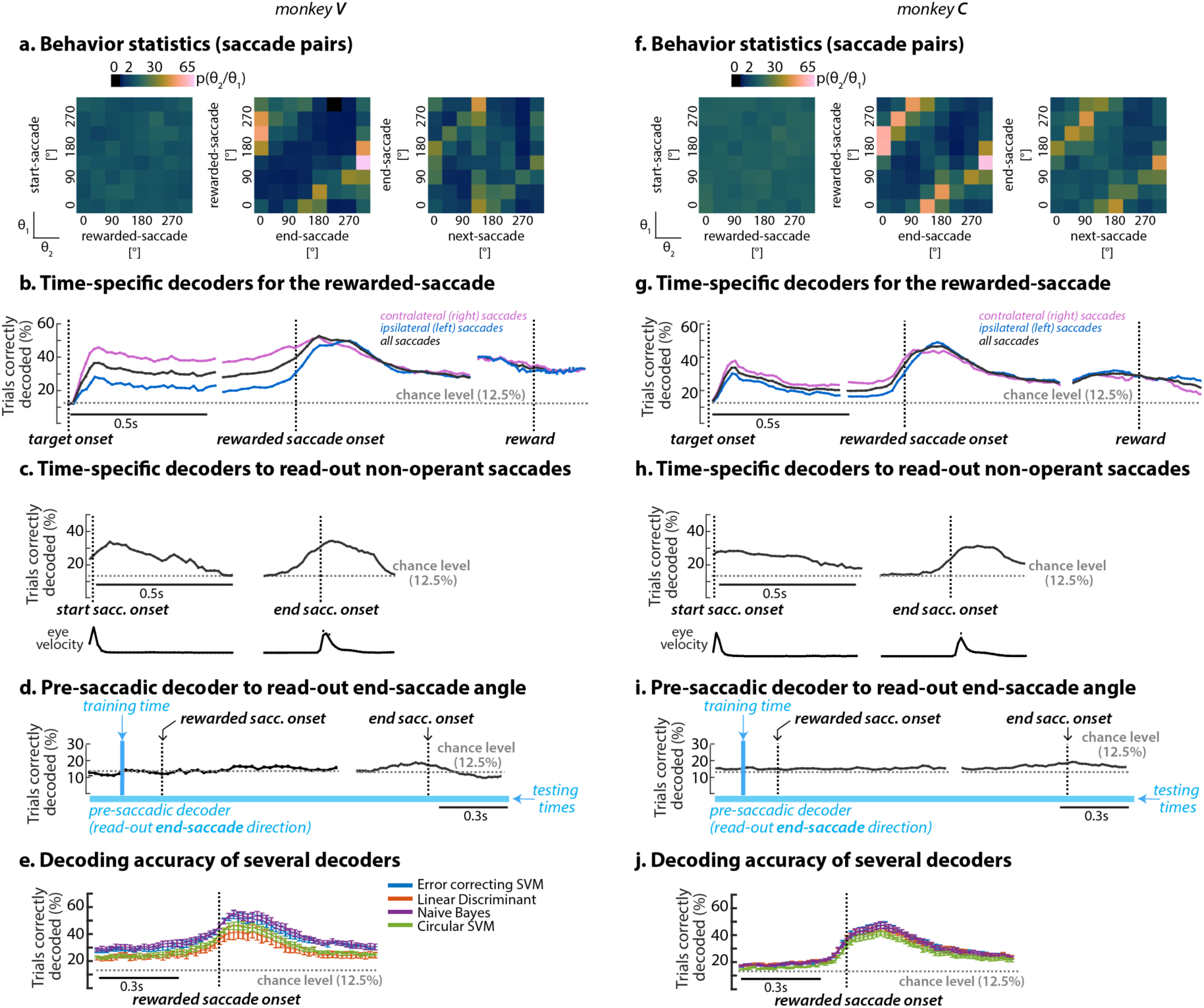
Saccade representations in monkeys V and C. **a-d**: Data for monkey V. **a.** Statistics of subsequent saccades, analogous to Suppl. Fig. 4. **b.** Time-specific decoding of the direction of the rewarded saccade, as in Figure 2a. **c.** Time-specific decoding of the direction of the start and end saccades, based on corresponding decoders trained on rewarded saccades. Analogous to Figure 2b. **d.** Decoding the direction of the end saccade, based on a pre-saccadic decoder trained on rewarded saccades. Analogous to Figure 3b. **e-h:** Analogous to a-d, but data from monkey C.

**Supplementary Figure 4.**
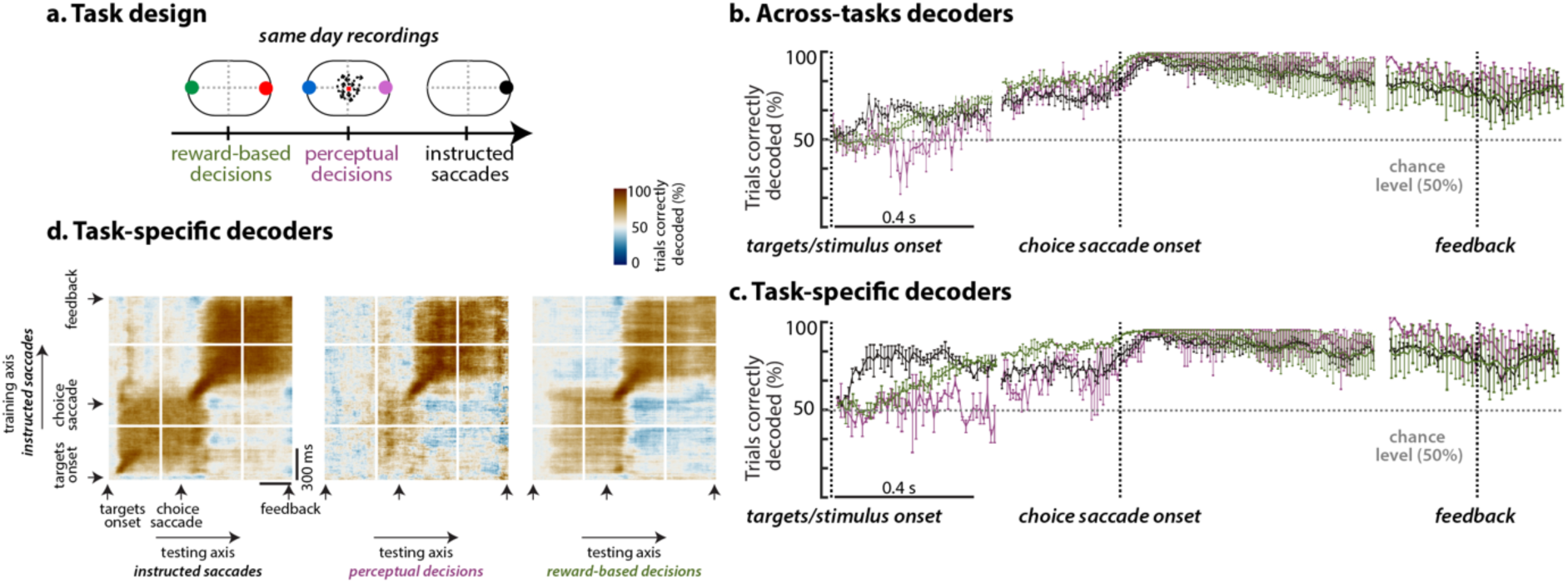
Different tasks, same patterns of activity (monkey V). Analogous to Fig. 4. Error bars indicate s.e.m. across sessions. (n=2)

**Supplementary Figure 5.**
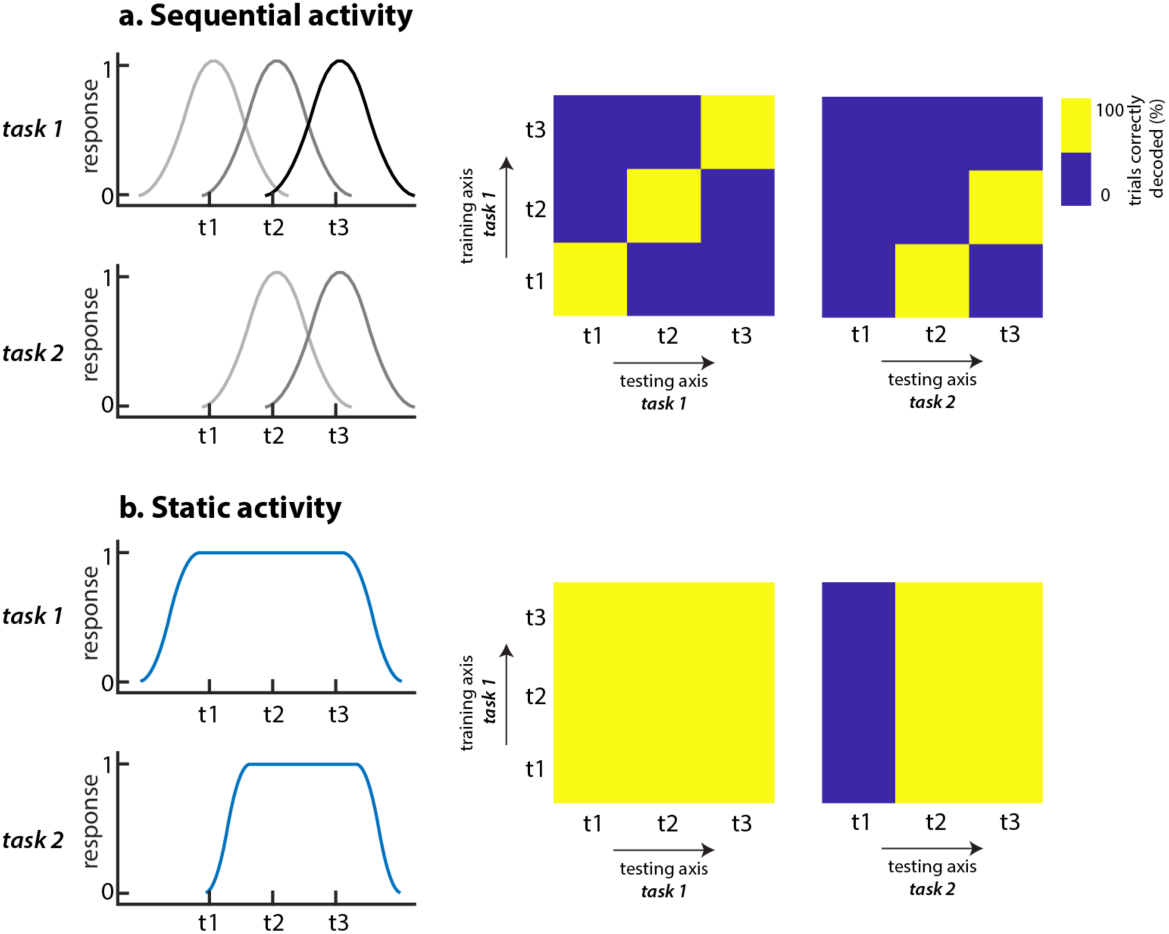
Hypotheses of choice-dynamics across tasks. Task-dependent temporal dynamics for two different choice representations. In both scenarios, we consider the case where choice activity in task 2 undergoes the same dynamics as in task 1, but its onset is delayed. a. A temporal sequence, where each pattern is active at a certain time. We apply the same analysis as in Fig. 4d, where we train time-dependent decoders on activity from task 1 and test them on activity from all times in task 1 (left panel) and task 2 (right panel). Training and testing on task 1 reveals a diagonal decoding matrix, with high values on the diagonal when the training and testing time coincide, and low values off-diagonal, where training and testing time differ. When testing the decoders on activity from task 2 (where choice activity is delayed), the decoding accuracies are high under the diagonal, matching the one time step delay set up in the cartoon of the time-courses. b. A static activity pattern. Training and testing on task 1 reveals a block decoding matrix, with high decoding values everywhere. Testing on activity from task 2 reveals a block structure starting at the second time step, again corresponding to the delay set up in the cartoon of the time-courses.

**Supplementary Figure 6.**
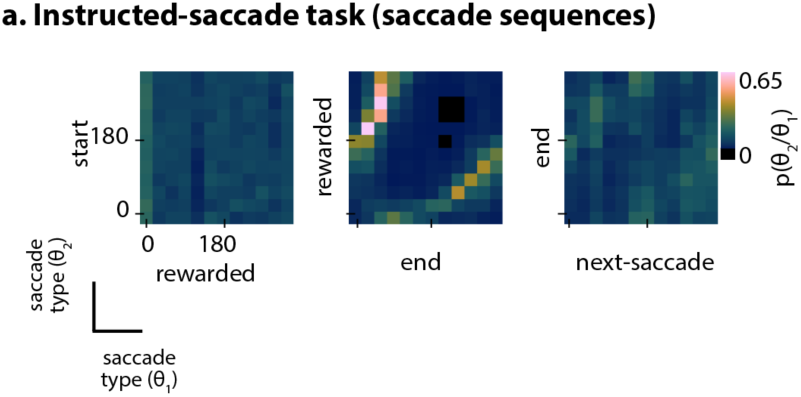
Saccade sequences in the instructed-saccade task for monkey T. Histogram of consecutive saccades, expressed as the distribution of directions for the second saccade (columns) conditional on the direction of the first saccade (rows), shown for the start and rewarded saccade (left); rewarded and end-saccade (middle); end and the following saccade (right).

**Supplementary Figure 7.**
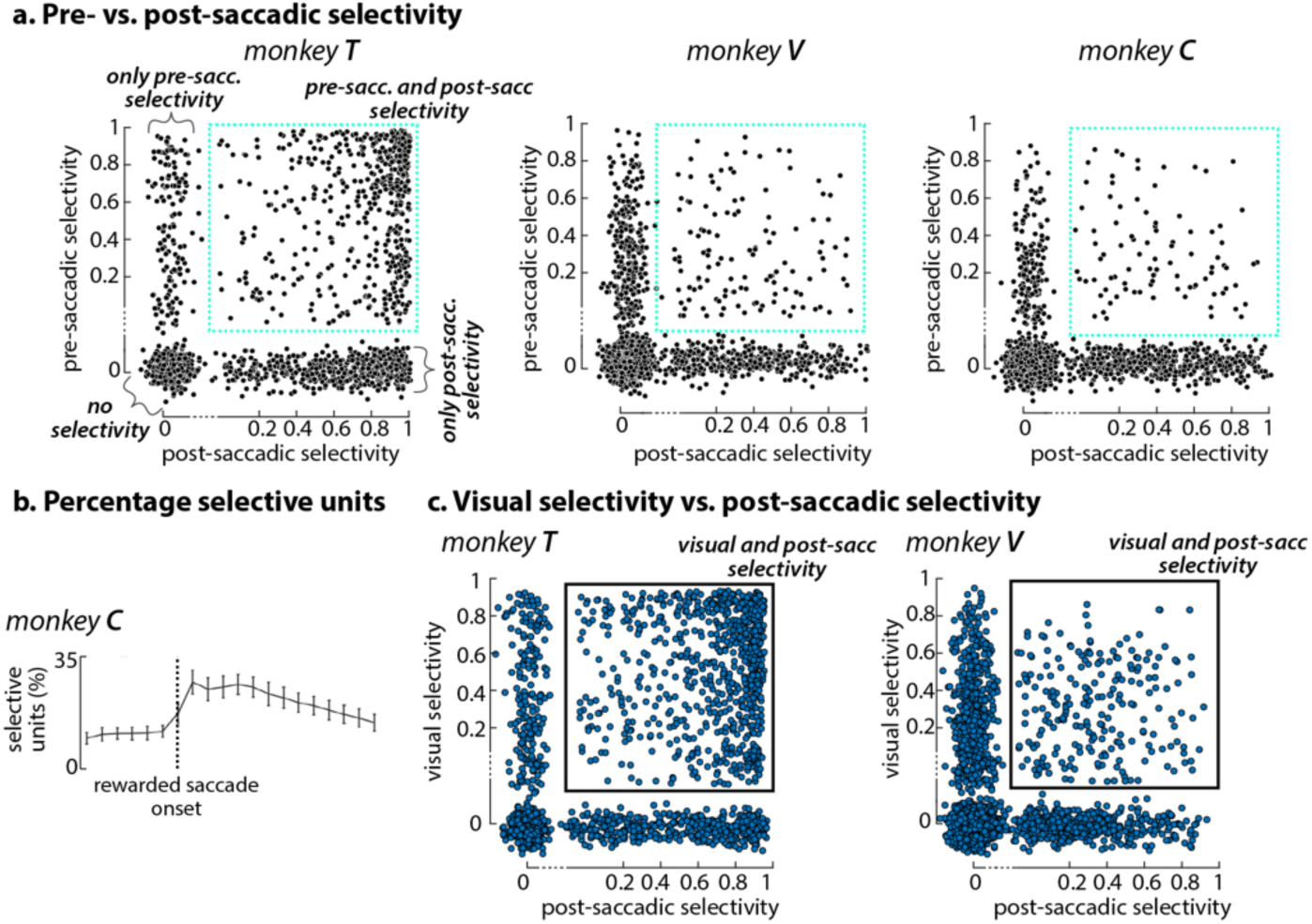
Unit direction-selectivity. **a.** Pre- vs post-saccadic selectivity. Pre-saccadic (-100ms, horizontal axis) vs. post-saccadic selectivity (+150ms, vertical axis), quantified as the goodness-of-fit (cross-validated r-squared) of the direction-tuning model at these times. Each point represents a unit. For each monkey, units from all sessions are displayed. Random jitter is added to units with “no selectivity” along both the vertical and horizontal axes; to units with only “pre-saccadic selectivity” along the horizontal axis; and to units with only “post-saccadic selectivity” along the vertical axis. Axes are linear and interrupted between 0 and 0.2 to be able to differentiate between units with “no selectivity” and units with only “pre-saccadic” or “post-saccadic” selectivity. **b.** Percentage of selective units computed on responses aligned to rewarded saccade onset for monkey V and C. Analogous to Fig. 5c, top row (right). **c.** Visual vs. post-saccadic selectivity. We consider a unit to have visual selectivity if it has direction-selectivity (cross-validated r-squared > 0 of the bell-shaped model in Fig. 5 applied to direction-averaged responses) at target onset. Here, we directly compare the visual selectivity at +100ms post target onset (vertical axis) against post-saccadic selectivity at +100ms post rewarded saccade onset (horizontal axis). Correlation of selectivity strength (defined as the r-squared) at these two times is for monkey T rho = 0.05, p-value = 0.2 and for monkey V rho = 0.02, p-value = 0.7. **d.** Cross-temporal selectivity for monkey V, analogous of Fig. 5c, bottom.

**Supplementary Figure 8.**
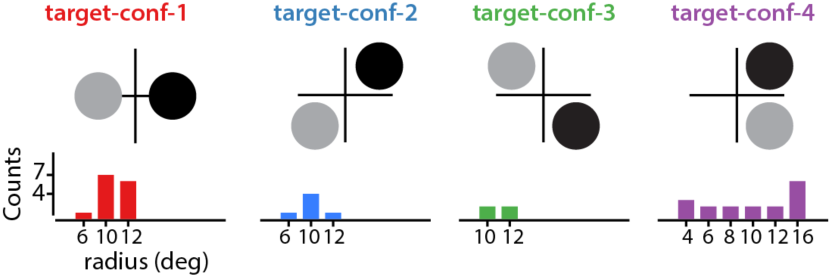
Task parameters in the learning task. **a**. The target locations during the reversal learning task can be in one of the five target configurations. Within each target configuration, either the radius may vary (classes 1-4) or the angular distance between the two targets (class 5). The bar plots indicate how many sessions there are for each target configuration and radius combination (data for monkey T).

**Supplementary Figure 9.**
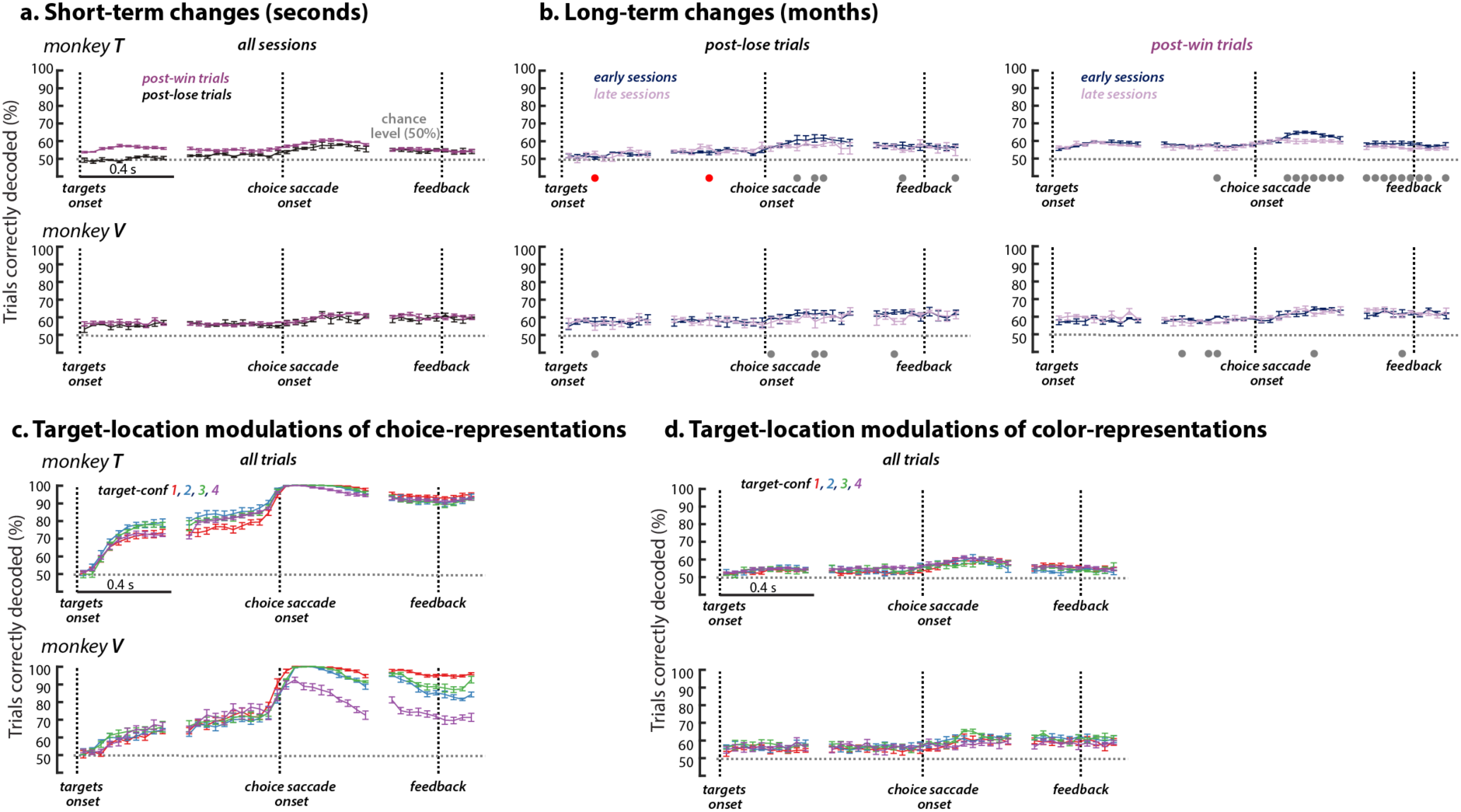
Color-representations during learning. **a,b** Analogous of Fig 8a,b. c, d Target-location effects on choice-representations **(c)** and color-representations (d). The target-placement (shown in Suppl. Fig. 8a) affected the decoding accuracy at different times of location decoders, but not of color decoders. We account for target-location (using one-hot encoding) when correlating decoding accuracy and training day (b for color and Fig. 8b for location)

**Supplementary Figure 10.**
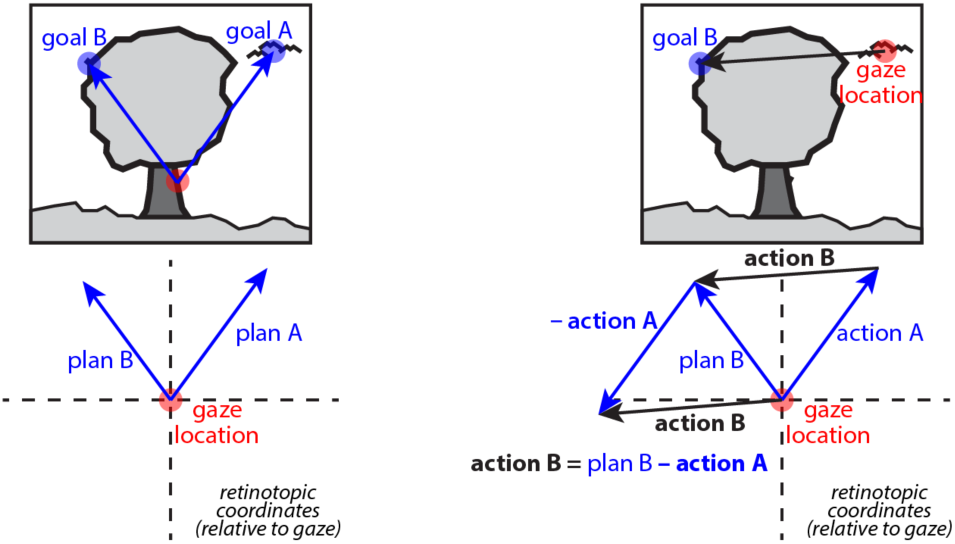
Vector subtraction mechanism for spatially accurate saccades. Diagram suggesting how a vector subtraction mechanism could be used to adjust sensory representations across saccades. Suppose the motor plan is to perform two consecutive saccades to two targets, goal A and goal B. Left panel: The motor plan is constructed while the gaze is at the bottom of the tree (red circle) and uses the retinal registration of the two targets, plan A and plan B. Right panel: The first saccade (action A) corresponds to plan A, and is thus a consonant-vector saccade. The second saccade is, on the other hand, a dissonant-vector saccade, because the movement vector does not correspond to the original retinal registration plan B. The movement-vector of the second saccade (action B) is obtained by subtracting the vector of the intervening (first) saccade from the retinal registration of the second target: action B = plan B - action A. See Fig. 13 in ^19^.

## 1 Experimental Procedures

We collected behavioral and neural data from three adult male rhesus monkeys: monkeys T (14 kg), V (11 kg) and C (13 kg). All surgical, behavioral, and animal-care procedures complied with National Institutes of Health guidelines and were approved by the Stanford University Institutional Animal Care and Use Committee. Prior to training, the monkeys were implanted with a stainless-steel head holder [1] and a scleral search coil for monitoring monocular eye position [2]. We used operant conditioning with liquid rewards to train the monkeys to perform a visually guided, delayed-saccade task; a two-alternative, forced-choice, motion discrimination task; and a two-alternative, forced-choice, non-spatial associative task. During training and experimental sessions, monkeys sat in a primate chair with their head restrained. Visual stimuli were presented on a cathode ray tube monitor controlled by a VSG graphics card (Cambridge Graphics, UK), at a frame rate of 120Hz, and viewed from a distance of 57 cm. Eye movements were monitored through the scleral eye coils (C-N-C Engineering, Seattle, WA). Behavioral control and data acquisition were managed by a computer running the REX software environment and QNX Software System’s (Ottawa, Canada) real-time operating system.

## 2 Behavioral tasks

### 2.1 Instructed saccade task

Monkeys were engaged in a visually-guided, delayed-saccade task, requiring them to perform a sequence of saccades and fixations on each trial to obtain a reward (Fig. 1a). A trial was initiated by a saccade to the fixation point, and subsequently the monkey was required to maintain fixation until the offset of the fixation point. At 0.6-0.8s after fixation onset, a saccade target was presented in the periphery (33 unique positions per experiment for monkey T and 24 for monkey V and C). The fixation cue disappeared after an interval of random duration following the target onset (0.7-1.2s) instructing the monkey to execute the saccade to the target. After the saccade, the monkey was again required to maintain fixation, this time on the target, for the duration of another random time interval (0.8-1.5s). At the end of this interval, the target disappeared, a reward was delivered, and the monkey was free to move the eyes. The three randomized intervals were drawn from uniform distribution.

Note that for monkey T possible targets were placed 30 degrees apart, but only 11 out of 12 ({0, 30, 60, 90, 120, 150, 180, 210, 240, 270, 300, 330} degrees) directions were used per experiment. Specifically, targets at 120 degrees were never present in 5 sessions; 300 degrees never appeared in 3 sessions and 210 degrees never appeared in one session. Each target direction could appear at one of three eccentricities or radii ({4, 8, 12}). For monkey V and monkey C, each recording session included 24 unique target locations (8 target directions placed at 45 degrees apart and 3 possible radii - 4, 8 and 12).

For the instructed saccade task, we analyzed neural recordings obtained when the monkeys were proficient at the task, i.e. there are no error trials and the direction of the rewarded saccade always refers to the target location. We analyzed a total of 9/10/10 experiments with 20,952/4751/8611 trials and 1706/2334/2095 single and multi-units from the three arrays in monkeys T/V/C.

### 2.2 Perceptual decision-making task (moving-dots)

Monkeys were engaged in a two-alternative, forced-choice motion discrimination task (Fig. 3d, e). The timing of task events was similar to the instructed saccade task (i.e. it included the random interval of target-fixation after the choice saccade). On each trial monkeys observed a noisy, random-dots motion stimulus presented through a circular aperture and had to report the prevalent direction of motion with a saccade towards one of two visual targets. Correct choices (e.g. a saccade to the right target for predominant rightward motion) were rewarded at the end of the target-fixation period. The strength of the motion stimulus (motion coherence) was set pseudo-randomly on each trial. For low motion coherences, the monkeys’ performance was close to chance level (50%), while for high coherences it was close to perfect (not shown). In this manuscript we only analyzed rewarded trials with high motion coherence stimulus.

### 2.3 Shifted workspace for the perceptual task

We used a modified version of the moving-dots task to investigate whether post-saccadic activity of the rewarded saccade is affected by the position of the eye (Fig. 3d). The timing of relevant task-events was analogous to that in the instructed saccade task, and included a target-fixation-period after the rewarded saccade (i.e. the choice saccade). Critically, each experiment in this task included trials from two “shifted” workspaces, whereby the location of the fixation point was shifted to the left from the midline in one workspace (relative to head-position), and to the right in the other (Fig. 3d, “left” and “right” workspaces). As a result, saccade direction and gaze-location of the rewarded saccade are somewhat decoupled—for example, the location corresponding to the center of the monitor could either be the target of a rightward or a leftward saccade (Fig. 3d, left vs. right workspace).

### 2.4 Non-spatial associative task

Monkeys were engaged in a two-alternative, forced-choice task that required them to track which of two targets (red or green) was being rewarded at any given time (Fig. 7a). Throughout the day, the reward contingencies switched repeatedly between two “contexts”: in the red context, only saccades to the red target were rewarded, and in the green context only saccades to the green target were rewarded. Because the timing of switches in reward contingencies was unpredictable, the optimal strategy is “win-stay-loose-switch”: if a given color was rewarded (“win”), the monkey should choose the same color again in the next trial (“stay”). Instead, after a choice that was not rewarded (“loose”) the monkeys should switch to the other color (“switch”).

### 2.5 Same recording day for perceptual task, instructed saccade and associative with 2 targets

On some recording days, monkey T (12 sessions) and monkey V (2 sessions) performed three tasks sequentially: the perceptual task (random-dots), the instructed saccade task where one target could appear in one of two locations and the non-spatial associative task (Fig. 4). Importantly, the target locations across the three tasks were identical, allowing the comparison of saccade-related activity across the different tasks.

## 3 Neural recordings

We recorded single and multi-unit neural signals with a chronically-implanted 10 by 10 array of electrodes (Cyberkinetics Neurotechnology Systems, Foxborough, MA; now Blackrock Microsystems). The inter-electrode spacing was 0.4 mm; electrodes were 1.5 mm long. Arrays were surgically implanted into the pre-arcuate gyrus[3,4]. We targeted the array to a region of prefrontal cortex between the posterior end of the principal sulcus, and the anterior bank of the arcuate sulcus, near the rostral zone of Brodmann’s area 8 (area 8Ar) in monkeys T and V. The arrays were implanted in the left hemisphere in both monkeys. The exact location of the array varied slightly across the two monkeys (Suppl. Fig. 1a), due to inter-animal variations in cortical vasculature and sulcal geometry that constrained the location of the array insertion site. In monkey C the array was placed between the superior branch of arcuate sulcus and dorsal bank of the principal sulcus, in the right hemisphere.

Array signals were amplified with respect to a common subdural ground, filtered and digitized using hardware and software from Cyberkinetics. For each of the 96 recording channels, ‘spikes’ from the entire duration of a recording session were sorted and clustered offline, based on a principal component analysis of voltage waveforms, using Plexon Offline Sorter (Plexon Inc., Dallas, Texas). This automated process returned a set of candidate action-potential classifications for each electrode that were subject to additional quality controls, including considerations of waveform shape, waveform reproducibility, inter-spike interval statistics, and the overall firing rate. For clusters returned by this postprocessing, both spike-waveform and spike-timing metrics fell within previously-reported ranges for array recordings[3].

Daily recordings yielded 100-200 single and multi-unit clusters distributed across the array. We do not differentiate between single-unit and multi-unit recordings, referring to both collectively as ‘units’. Therefore, we also do not draw conclusions in this study that depend on the distinction between single and multi-unit responses. Neural responses in the instructed saccade task were recorded over a total of 9, 10, 10 experiments in monkeys monkey T, monkey V and monkey C, for a total of 20,905, 4751 and 8611 trials.

## 4 Analysis of eye movement data

### 4.1 Saccade extraction

We used a non-parametric data-driven method for classifying eye fixations and saccades that automatically adapts itself to the task statistics[5]. The method is built on the assumption that the eye reaches higher speeds during saccades than during fixations, and that there are fewer peaks in speed due to saccades than due to fixations. Using these observations about the statistics of eye-behavior, the method derives an optimum speed threshold that best separates the speed distribution of saccades from the speed distribution of fixations and instrumental noise.

### 4.2 Saccade types

We analyze neural activity related to different types of saccades, i.e. the instructed and freely initiated saccades occurring before, during, and after each trial (Fig. 1b, c). We refer to the initial saccade to the fixation point as the start saccade, the saccade to the target as the rewarded saccade, and the saccade away from the target after reward delivery as the end saccade. The start saccade is therefore visually-guided and non-rewarded; the rewarded saccade is visually-guided and rewarded; and the end saccade is free and non-rewarded. Monkeys initiate the end saccade when there is nothing on the screen. The saccade durations are 30+-30ms, 40+-10ms and 140+-80ms for monkey T, for the start, rewarded and end saccades respectively.

In the instructed saccade task, in approximately 45% (monkey T) and 33% (monkey V) trials the monkeys were already at fixation point when the new trial started, thus there are fewer start saccades than end saccades.

## 5 Analysis of neurophysiology data

Throughout the paper, we consider neural responses occurring during four distinct, largely non-overlapping trial epochs. We refer to the first randomized time interval, following the start saccade, as the first central-fixation-period (i.e. fixation on the fixation point, 0.6-0.8s); the second randomized interval, preceding the rewarded saccade, as the second central-fixation-period (0.7-1.2s); and the last randomized interval, preceding the reward, as the target-fixation-period (i.e. fixation on the target, 0.8-1.5s). Lastly, we analyze the times around the end saccade, whose onset is after reward delivery. Notably, the onset of the end saccade does not coincide with the time of reward delivery on every single trial - on some trials monkeys initiate the end-saccade immediately after reward and on some trials monkeys continue fixating the location where the target was present, on a few trials for intervals as long as 600ms.

### 5.1 Unit-specific direction selectivity

#### 5.1.1 Pre-processing condition-averaged responses

We bin activity in 50ms non-overlapping bins and we normalize the unit responses using z-scoring:

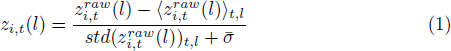

where 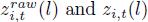 are the raw firing rate and *z*-scored responses, respectively, of unit i at time t and on trial 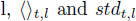 indicate the mean and standard deviation across times and trials, and 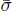 is a constant defined as the median of the standard deviation across all units in a session. The *z*-scoring deemphasizes the contribution to the population response of units with very high firing rates (typically multi-unit activity), while the constant term ensures that units with very small firing rates are not over-emphasized. For the unit-level analysis, we do not apply any other temporal smoothing to the responses.

We defined condition-averaged responses *f_i,t,c_* for each unit by averaging the normalized time-varying firing rates across all trials belonging to a given condition c (Fig. 5a). For the instructed saccade task, we define each condition by the saccade direction (11 conditions for monkey T, 8 conditions for monkey V and monkey C).

The condition-averaged responses were de-noised using Singular Vector Decomposition (SVD). We concatenated the condition-averaged responses *f_i,t,θ_* across all recording sessions with the same conditions in a *N_unit_* x *(N_condition_* • *T*) matrix, where *N_unit_* is the total number of units, *N_condition_* is the total number of conditions, and T is the number of bins. The left singular-vectors of this data matrix are vectors v_a_ of length *N_unit_*, indexed by a, ordered from the singular-vector explaining the most variance to the one explaining the least. We use the first *N_svd_* singular-vectors to define a de-noising matrix D of size *N_unit_* x *N_unit_* :

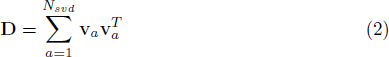

We used this matrix to de-noise the condition-averaged responses by projecting them into the sub-space spanned by the first *N_svd_* singular-vectors:

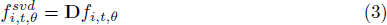

We use the de-noised condition-averaged responses 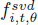 to determine the unitspecific optimal direction, i.e. the condition that elicits the highest responses. From now on *f_i,t,θ_* will refer to the de-noised responses.

#### 5.1.2 Bell-shaped model of direction selectivity

We estimated, for each unit, the saccade-location that elicits the highest response at each time by fitting a descriptive function[6] to the normalized timevarying condition-averaged responses (Fig. 5b):

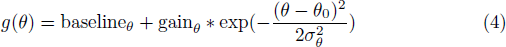

where *θ_0_* is the preferred saccade direction, *σ_θ_* determines the tuning width and gain*_θ_* determines the modulation depth of the tuning curve.

We fitted the parameters of these models separately for each unit to averaged responses grouped by saccade-direction within the epoch [0, 0.7]s after target onset and [-0.3, 0.5]s around saccade initiation, in 50ms non-overlapping bins. The models are fit by minimizing the summed square error across the respective conditions between the model predictions and the corresponding condition-averaged response.

#### 5.1.3 Goodness-of-fit

We validated the 1-D bell-shaped models by computing a coefficient of determination R^2^ (Fig. 5b) value from the measured condition-averaged response *f_i,t,θ_* and the model’s reconstruction 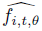, based on comparing the variability of the estimation errors with the variability of the original neural responses.

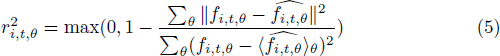

Model parameters were found from condition-averages computed on a subset of trials (training set) and validated on condition-averages computed on a different, non-overlapping subset of trials (testing set). All units that had a coefficient of determination different than 0 were considered selective (Fig. 5c, top row). A coefficient of determination equal to 0 indicates that the condition-averaged response is better described by the averaged response across all conditions 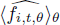.

#### 5.1.4 Cross-temporal selectivity measure

We quantified the percentage of selective units at different time-pairs (*t_m_, t_n_*) (Fig. 5c, bottom row):

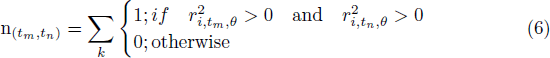

where i is unit index.

To assess the significance of each n(*t_m_,t_n_*), we shuffled the unit-order independently at *t_m_* and *t_n_* and re-computed the number of units that were selective at both times. We repeated this procedure 1000 times and compared the measured n(*t_m_,t_n_*) to the 95th percentile of this distribution.

### 5.2 Population Decoding

For the population-level analysis, we compute binned spike counts in 100ms overlapping bins. Chance level of decoding analyses is computed using 11 classes (9%) for monkey T and 8 classes (12.5%) for monkey V and monkey C. We quantified the relation between single-trial normalized population responses and the saccade direction using high-dimensional decoders suited for multi-class problems (Fig. 2 for monkey T, Suppl. Fig. 3b, g for monkey V and C). To ensure our results do not depend on the choice of the decoder, we used several types of decoders (Suppl. Fig. 2e for monkey T, Suppl. Fig. 3e, j for monkey V and C). Specifically, we used MATLAB built-in classifiers: Linear discriminant analysis (fitcdiscr), Naive Bayes (fitcnb) and Error-correcting SVM (fitcecoc), as well as a customized classifier (Circular-SVM).

The Circular-SVM was proposed by Graf et al.[7] and builds on the Naïve Bayes model. Knowing that the topography of the neural responses is circular, it learns the pooling weight W, i.e. how each unit influences the classifier’s prediction, in a model-free way, directly from the neural data. We describe the method briefly, for more details see[7].

Discrimination between two saccade directions *θ*_1_ and *θ*_2_ is done using the sign of the Support-Vector Machine (SVM) decision function:

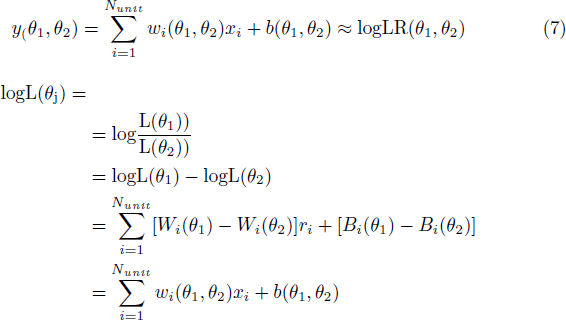

The SVM decision function is used as a local linear approximation of the difference between the log-likelihood evaluated at two saccade directions. The entire log-likelihood function is reconstructed by computing the cumulative sum of the empirical log-likelihood ratios of adjacent directions:

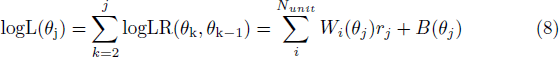

with log(*θ*_1_) = 0.

Some pairs of neighboring directions are better separated than others. We modified the original version of the method such that the discriminability of a saccade-direction would only depend on how well it is separated from its two immediate neighboring directions, and not on how well separated are any other two neighboring directions. To compute an unbiased log-likelihood, each angle *θ_j_* takes turn in being the reference log(*θ_j_*) = 0. In this manner, we average out the cumulated-error.

#### 5.2.1 Decoding saccade direction of the start, rewarded and end saccades

Rewarded saccade: We study the relationship between the population responses and the rewarded saccade direction through cross-validated high-dimensional decoders (Fig. 2a for monkey T, Suppl. Fig. 3b, g for monkey V and C).

Start and end saccade: We apply the same decoders we identified for the rewarded saccade to responses aligned to the start and end saccade (Fig. 2b for monkey T, Suppl. Fig. 3c, h for monkey V and C). Training a new set of decoders on responses aligned to the end saccade resulted on similar cross-validated accuracies when used to read-out the end saccade (results not shown).

#### 5.2.2 Time-specific decoding

Decoders are trained and tested on time-specific responses using 10-fold cross-validation.

#### 5.2.3 Cross-temporal decoding

Decoders are tested on responses outside their training time-window (Fig. 4d). A decoding matrix *T* x *T* contains the cross-validated decoding accuracy of T time-specific decoders tested on T time-specific population-responses. The diagonal of this decoding matrix is the time-specific decoding accuracy. All decoders are cross-validated, i.e. that even though the decoders are trained at one time and tested at another time, there is no overlap between the train and test trials. This analysis shows how each of the time-specific mappings generalize across responses at other times in the trial.

### 5.3 Post-saccadic activity is not pre-saccadic activity for the next saccade

One possible interpretation of post-saccadic activity is that it encodes the planning of the next saccade (Fig. 3a-c). To test this hypothesis, we decoded the direction of the end saccade from activity preceding the end saccade and from activity during the target-fixation-period (Fig. 3b). Ruling out this hypothesis is very challenging because the behaviour of the monkeys is biased - very often the end saccade is back to the fixation point.

To study this, we used a pre-trained pre-saccadic decoder. Specifically, we used a decoder trained to decode the rewarded-saccade during the pre-saccadic epoch (t : t + Δt where t = -150ms and = -50ms) to decode the saccade direction across the target-fixation and up until the onset of the end-saccade. Importantly, we use the decoder to read out the direction of the end saccade, not of the rewarded saccade and we evaluate the accuracy of the read-outs separately for trials from a single direction of the rewarded saccade. We focus on rewarded saccades to the contralateral hemifield, which are followed by end saccades in many different directions and are thus well-suited to test the decoder (rewarded saccades towards 0, 30 and 60 degrees in Fig. 3a, left panel).

Figure 3b shows that post-saccadic activity following the rewarded saccade does not contain preparatory activity for the end saccade, when these behavioural correlations are “subtracted” (see histogram of balanced conditions in Figure 3a, right panel), but does contain information about the rewarded saccade. Importantly, the decoding accuracies are computed from the same trials in both cases. Note that it is still possible that preparatory activity of the end saccade would exist along another read-out, one that is different from the pre-saccadic read-out of the rewarded saccade. Even so, this result shows that the inverted tuning of pre-saccadic activity after saccade execution is not a consequence of the next saccade the monkey will perform.

### 5.4 Prospective and retrospective representations have different task-selectivity

On some recording days monkeys performed three tasks sequentially: (1) the perceptual decision-making task, where the monkeys had to choose between two targets based on sensory information; (2) the instructed-saccade task, where only one peripheral target was presented on each trial; (3) the non-spatial associative task, where monkeys had to choose between two targets based on information from the previous trial. Targets were placed at identical locations across the three tasks, allowing us to study how task-context modulates responses.

We analyzed responses aligned to target (one target in instructed saccade task, two targets of different colors in the associative task and dots onset in the perceptual task) and saccade onset. We identify choice-decoders that best separate the population responses due to monkey’s choices (leftward or rightward) across the three tasks.

Note that because trials within the three tasks are not intermingled, but come in sequential blocks, we corrected the single-trial spike counts of any potential population-level drift in the baseline firing rates:

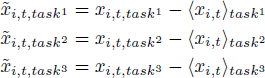

The decoding analyses was performed on the normalized responses.

### 5.5 Post-saccadic activity does not encode the momentary gaze location

We addressed the question whether post-saccadic activity is better explained by saccade-covariates or eye-position-covariates in a modified version of the perceptual decision-making task, in which the monkeys were presented with two workspace configurations in a blocked design (Fig. 3d, e). The task required the monkeys to discriminate the dominant movement of moving dots in two “workspaces” that were retinotopically identical, but horizontally (or vertically) shifted along the monkey’s line of sight, such that the physical location of one target (T1) in one block was identical to the physical location of the other target (T2) in the other block.

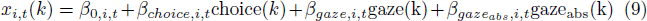

where *x_i,t_* (*k*) is the z-scored response of unit i at time t and on trial k, choice(k) is the monkey’s choice on trial k (+1 for choice 1 and -1 for choice 2), gaze(k) is the target-location on trial k (for two sessions the workspace is shifted along the horizontal axis gaze = gaze**_x_** = {-1, 0, 1} and gaze**_y_** = 0; and for two sessions the workspace is shifted along the vertical axis gaze = gaze_y_ = {-1, 0, 1} and gaze**_x_** = 0, gaze_abs_(k) is the absolute value of gaze(k). We introduced gaze_abs_ (k) to capture a potential non-linear relation between neural responses and gaze. We focused on three time points in the post-saccadic epoch: early (+50ms), middle (+200ms) and late (+400ms).

Because trials within the two retinotopically-identical sessions, workspace_1_ and workspace_2_, are not intermingled, but come in sequential blocks, we corrected the single-trial spike counts of any potential population-level drift in the baseline firing rates:

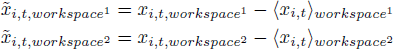

We identified the regression coefficients 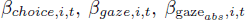 through 10-fold cross-validation for each unit separately. We next quantified the saccade-related and gaze-related contributions of each unit through a measure of variance explained on the test trials:

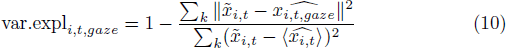

where

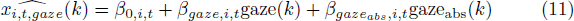

Similarly, for saccade-related activity:

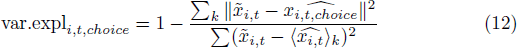

### where

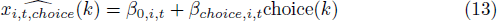

## 6 Analysis of behavioral data in the non-spatial associative task

We characterized fast learning with logistic regression models fit to the behavior in a single session. We separately modeled the influence of the (task relevant) target color and the (task irrelevant) target location on the monkeys’ choices (Fig. 7c, circles and squares) and their interaction with previous outcome (win or lose, x- and y-axes).

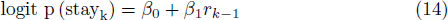

where logit p (stay_k_) denotes the probability of choosing on trial k the same choice (same colour for the optimal strategy and same location for the suboptimal) as in the previous trial k-1 and r*_k_*_-_**_1_** is 1 when the outcome of the previous trial was a reward and 0 when it was not rewarded.

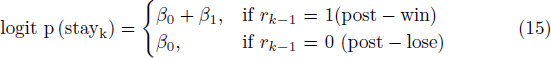

Then, we compute logit p (stay_k_) in post-win trials:

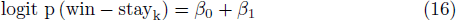

and logit p (switch_k_) in post-lose trials:

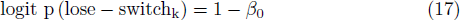

In Fig. 7c we subtract from these probabilities the estimated probabilities of random behaviour and obtain ΔP(lose — switch) and ΔP(win — stay). For the optimal model (colour), we simulate random colour-choices . Fitting the logistic regression in Eq. 14, we estimate p(win — stay^color^) =0.5 and p(lose — switch^color^) = 0.5. Then, for the colour-model,

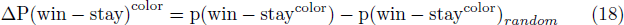

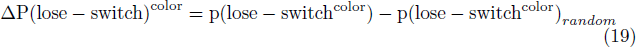

The relation between space and color was pseudo-randomized in some sessions and this resulting deviation from perfect randomness lead to apparent biases in the behavior that we seek to remove. Using such relative probabilities allowed us to compensate for any potential biases in choices due to lack of complete randomization (e.g. because of the limited number of switches in a trial, and the fact that timing of transitions is pseudo-randomized, rather than being completely random). Therefore, for the location-model, we corrected the estimated probabilities of the monkey’s behaviour by (1) using their observed colour-choices; (2) converting the colour-choice into a location-choice, based on the experimentally-set colour-location association of each trial; (3) estimating p(lose - switch^location^)_random_ and p(win - stay^location^)_random_.

## 7 Correlation of behavioral data and neural data in the non-spatial associative task

We studied how slow learning shapes the neural responses by correlating timespecific decoding accuracies with different behavioral variables. Notably, we used partial correlation to remove modulations due to target configuration (Suppl. Fig. 9c, d). Target configuration was included through one-hot encoding. Behavioral variables we considered are: training day, first half vs second half training day (1 for first half and 2 for second half) and the modeled probabilities of the task relevant model. The results were quantitively similar. The p-values in Fig. 8b and Suppl. Fig. 9b are for the partial correlation with first half vs second half training day. P-value is computed as the number of shuffled partial correlations that exceed the empirical partial correlation. Shuffled partial correlations were computed by correlating behavioral variable to 1000 random permutations of decoding accuracies.

## References

1. Mansouri, F., Tanaka K & Buckley MJ. Conflict-induced behavioural adjustment: a clue to the executive functions of the prefrontal cortex - PubMed. Nat Rev Neurosci 10, 141–152 (2009).

2. Fuster, J. M. The prefrontal cortex. (Academic Press/Elsevier, 2008).

3. Miller, E. K. & Cohen, J. D. An Integrative Theory of Prefrontal Cortex Function. Annu. Rev. Neurosci. 24, 167–202 (2001).

4. Duncan, J. & Miller, E. K. Cognitive Focus through Adaptive Neural Coding in the Primate Prefrontal Cortex. in Principles of Frontal Lobe Function 278–291 (Oxford University Press, 2002). doi:10.1093/acprof:oso/9780195134971.003.0018

5. JM, F. The prefrontal cortex--an update: time is of the essence. Neuron 30, 319–333 (2001).

6. Duncan, J. An adaptive coding model of neural function in prefrontal cortex | Nature Reviews Neuroscience. Nat. Rev. Neurosci. 2, 820–829 (2001).

7. Mante, V., Sussillo, D., Shenoy, K. V. & Newsome, W. T. Context-dependent computation by recurrent dynamics in prefrontal cortex. Nature 503, 78–84 (2013).

8. Funahashi, S., Bruce, C. J. & Goldman-Rakic, P. S. Neuronal activity related to saccadic eye movements in the monkey’s dorsolateral prefrontal cortex. J. Neurophysiol. 65, 1464–1483 (1991).

9. Funahashi, S., Bruce, C. J. & Goldman-Rakic, P. S. Mnemonic coding of visual space in the monkey’s dorsolateral prefrontal cortex. J. Neurophysiol. 61, 331–349 (1989).

10. Fuster, J. M. & Alexander, G. E. Neuron activity related to short-term memory. Science (80-.). 173, 652–654 (1971).

11. Bruce, C. J. & Goldberg, M. E. Primate frontal eye fields. I. Single neurons discharging before saccades. J. Neurophysiol. 53, 603–35 (1985).

12. Asaad, W. F., Rainer, G. & Miller, E. K. Neural Activity in the Primate Prefrontal Cortex during Associative Learning. Neuron 21, 1399–1407 (1998).

13. Fuster, J. M., Bauer, R. H. & Jervey, J. P. Cellular discharge in the dorsolateral prefrontal cortex of the monkey in cognitive tasks. Exp. Neurol. 77, 679–94 (1982).

14. Rainer, G., Asaad, W. F. & Miller, E. K. Selective representation of relevant information by neurons in the primate prefrontal cortex. Nature 393, 577–9 (1998).

15. Miller, E. K., Erickson, C. A. & Desimone, R. Neural mechanisms of visual working memory in prefrontal cortex of the macaque. J. Neurosci. 16, 5154–67 (1996).

16. Romo, R., Brody, C. D., Hernández, A. & Lemus, L. Neuronal correlates of parametric working memory in the prefrontal cortex. Nature 399, 470–473 (1999).

17. Rigotti, M. et al. The importance of mixed selectivity in complex cognitive tasks. Nature 497, 585–590 (2013).

18. Bizzi, E. Discharge of frontal eye field neurons during saccadic and following eye movements in unanesthetized monkeys. Exp. Brain Res. 6, 69–80 (1968).

19. Goldberg, M. E. & Bruce, C. J. Primate frontal eye fields. III. Maintenance of a spatially accurate saccade signal. J. Neurophysiol. 64, 489–508 (1990).

20. Tsujimoto, S. & Sawaguchi, T. Context-dependent representation of response-outcome in monkey prefrontal neurons - PubMed. Cereb Cortex 15, 888–898 (2005).

21. Barraclough, D. J., Conroy, M. L. & Lee, D. Prefrontal cortex and decision making in a mixed-strategy game. Nat. Neurosci. 7, (2004).

22. Seo, H., Barraclough, D. J. & Lee, D. Dynamic Signals Related to Choices and Outcomes in the Dorsolateral Prefrontal Cortex. Cereb. Cortex 17, i110–i117 (2007).

23. Tsutsui, K. I., Grabenhorst, F., Kobayashi, S. & Schultz, W. A dynamic code for economic object valuation in prefrontal cortex neurons. Nat. Commun. 7, 1–16 (2016).

24. Donahue, C. H. & Lee, D. Dynamic routing of task-relevant signals for decision making in dorsolateral prefrontal cortex. Nat. Neurosci. 18, 295–301 (2015).

25. Olson, C. R., Musil, S. Y. & Goldberg, M. E. Single neurons in posterior cingulate cortex of behaving macaque: eye movement signals. J. Neurophysiol. 3285–3300 (1996).

26. Umeno, M. M. & Goldberg, M. E. Spatial Processing in the Monkey Frontal Eye Field. II. Memory Responses. J. Neurophysiol. 86, 2344–2352 (2001).

27. Tsujimoto, S., Genovesio, A. & Wise, S. P. Monkey orbitofrontal cortex encodes response choices near feedback time. J. Neurosci. 29, (2009).

28. Tsujimoto, S., Genovesio, A. & Wise, S. P. Evaluating self-generated decisions in frontal pole cortex of monkeys. Nat. Neurosci. 2009 131 13, 120–126 (2010).

29. Tsujimoto, S. & Sawaguchi, T. Neuronal representation of response-outcome in the primate prefrontal cortex - PubMed. Cereb Cortex. 14, 47–55 (2004).

30. Funahashi, S. Saccade-related activity in the prefrontal cortex: its role in eye movement control and cognitive functions. Front. Integr. Neurosci. 8, 54 (2014).

31. Xu, B. Y., Karachi, C. & Goldberg, M. E. The Postsaccadic Unreliability of Gain Fields Renders It Unlikely that the Motor System Can Use Them to Calculate Target Position in Space. Neuron 76, 1201–1209 (2012).

32. Khanna, S. B., Scott, J. A. & Smith, M. A. Dynamic shifts of visual and saccadic signals in prefrontal cortical regions 8Ar and FEF. J. Neurophysiol. 124, 1774–1791 (2020).

33. Kiani, R. et al. Natural Grouping of Neural Responses Reveals Spatially Segregated Clusters in Prearcuate Cortex. Neuron 85, 1359–1373 (2015).

34. Pasupathy, A. & Miller, E. K. Different time courses of learning-related activity in the prefrontal cortex and striatum. Nature 433, 873–876 (2005).

35. Law, C.-T. & Gold, J. I. Neural correlates of perceptual learning in a sensory-motor, but not a sensory, cortical area | Nature Neuroscience. Nat. Neurosci. 11, 505–513 (2008).

36. Sommer, M. A. & Wurtz, R. H. Composition and topographic organization of signals sent from the frontal eye field to the superior colliculus. J. Neurophysiol. 83, 1979– 2001 (2000).

37. Takeda, K. & Funahashi, S. Prefrontal Task-Related Activity Representing Visual Cue Location or Saccade Direction in Spatial Working Memory Tasks. (2002). doi:10.1152/jn.00249.2001

38. Bruce, C. J., Goldberg, M. E., Bushnell, M. C. & Stanton, G. B. Primate frontal eye fields. II. Physiological and anatomical correlates of electrically evoked eye movements. J. Neurophysiol. 54, 714–734 (1985).

39. Funahashi, S., Chafee, M. & Goldman-Rakic, P. Prefrontal neuronal activity in rhesus monkeys performing a delayed anti-saccade task. Nature 365, 753–756 (1993).

40. Bullock, K. R., Pieper, F., Sachs, A. J. & Martinez-Trujillo, J. C. Visual and presaccadic activity in area 8Ar of the macaque monkey lateral prefrontal cortex. J. Neurophysiol. 118, 15–28 (2017).

41. Murray, J. D. et al. Stable population coding for working memory coexists with heterogeneous neural dynamics in prefrontal cortex. Proc. Natl. Acad. Sci. U. S. A. 114, 394–399 (2017).

42. Cueva, C. J. et al. Low-dimensional dynamics for working memory and time encoding. Proc. Natl. Acad. Sci. U. S. A. 117, 23021–23032 (2020).

43. Khaki, M., Luna, R., Mortazavi, N., Sachs, A. & Martinez-Trujillo, J. Using classification-based decoding to analyze the Spatiotopic and Retinotopic memory representations in primates. J. Vis. 21, 2868–2868 (2021).

44. Luna, R. et al. Small neuronal ensembles of primate lateral prefrontal cortex encode spatial working memory in two reference frames. J. Vis. 21, 2858–2858 (2021).

45. Almeida, R. L., Roussy, M. P., Sachs, A., Treue, S. & Martinez-Trujillo, J. C. Neuronal ensembles of primate Lateral Prefrontal Cortex encode spatial working memory in different frames of reference. J. Vis. 20, 1753–1753 (2020).

46. Stokes, M. G. et al. Dynamic coding for cognitive control in prefrontal cortex. Neuron 78, 364–75 (2013).

47. Andersen, R. & Mountcastle, V. The influence of the angle of gaze upon the excitability of the light-sensitive neurons of the posterior parietal cortex. J. Neurosci. 3, 532–548 (1983).

48. Brotchie, P. R., Andersen, R. A., Snyder, L. H. & Goodman, S. J. Head position signals used by parietal neurons to encode locations of visual stimuli. Nature 375, 232–235 (1995).

49. Andersen, R. A., Essick, G. K. & Siegel, R. M. Encoding of spatial location by posterior parietal neurons. Science (80-.). 230, 456–458 (1985).

50. RA, A., RM, B., S, B., JW, G. & L, F. Eye position effects on visual, memory, and saccade-related activity in areas LIP and 7a of macaque. J. Neurosci. 10, 1176–1196 (1990).

51. Salinas, E. & Abbott, L. F. Chapter 11 Coordinate transformations in the visual system: how to generate gain fields and what to compute with them. Prog. Brain Res. 130, 175–190 (2001).

52. Meister, M. L. R. & Buffalo, E. A. Neurons in Primate Entorhinal Cortex Represent Gaze Position in Multiple Spatial Reference Frames. J. Neurosci. 38, 2430–2441 (2018).

53. Kim, S., Hwang, J. & Lee, D. Prefrontal Coding of Temporally Discounted Values during Intertemporal Choice. Neuron 59, 161–172 (2008).

54. Cohen, J. D., McClure, S. M. & Yu, A. J. Should I stay or should I go? How the human brain manages the trade-off between exploitation and exploration. Philos. Trans. R. Soc. B Biol. Sci. 362, 933–942 (2007).

55. Domjan, M. & Grau, J. W. The principles of learning and behavior. (Cengage Learning, 2015).

56. Sugrue, L. P., Corrado, G. S. & Newsome, W. T. Matching behavior and the representation of value in the parietal cortex. Science 304, 1782–7 (2004).

57. Parthasarathy, A. et al. Mixed selectivity morphs population codes in prefrontal cortex. Nat. Neurosci. 20, 1770–1779 (2017).

58. Spaak, E., Watanabe, K., Funahashi, S. & Stokes, M. G. Stable and dynamic coding for working memory in primate prefrontal cortex. J. Neurosci. 37, 6503–6516 (2017).

59. Raposo, D., Kaufman, M. T. & Churchland, A. K. A category-free neural population supports evolving demands during decision-making. (2014). doi:10.1038/nn.3865

60. Kim, J.-N. & Shadlen, M. N. Neural correlates of a decision in the dorsolateral prefrontal cortex of the macaque. Nat. Neurosci. 2, 176–185 (1999).

61. Petrides, M. Deficits in non-spatial conditional associative learning after periarcuate lesions in the monkey.

62. Marmor, O., Pollak, Y., Doron, C., Helmchen, F. & Gilad, A. History information emerges in the cortex during learning. Elife 12, (2023).

63. Kiani, R., Cueva, C. J., Reppas, J. B. & Newsome, W. T. Dynamics of Neural Population Responses in Prefrontal Cortex Indicate Changes of Mind on Single Trials. Curr. Biol. 24, 1542–1547 (2014).

64. Purcell, B. A. & Kiani, R. Neural Mechanisms of Post-error Adjustments of Decision Policy in Parietal Cortex. Neuron 89, 658–71 (2016).

65. Rabbitt, P. & Rodgers, B. What does a man do after he makes an error? an analysis of response programming: Quarterly Journal of Experimental Psychology: Vol 29, No 4. *Q. J. Exp*. Psych. 29, 727–743 (1977).

66. Schall, J. D. Visuomotor Areas of the Frontal Lobe. 527–638 (1997). doi:10.1007/978-1-4757-9625-4_13

67. Libby, A. & Buschman, T. J. Rotational dynamics reduce interference between sensory and memory representations. Nat. Neurosci. 24, 715–726 (2021).

68. Steel, A., Silson, E. H., Garcia, B. D. & Robertson, C. E. nature neuroscience A retinotopic code structures the interaction between perception and memory systems. Nat. Neurosci. | 27, 339–347 (2024).

69. Bizzi, E. & Schiller, P. H. Single unit activity in the frontal eye fields of unanesthetized monkeys during eye and head movement. Exp. Brain Res. 10, 151–158 (1970).

70. Sendhilnathan, N., Basu, D., Goldberg, M. E., Schall, J. D. & Murthy, A. Neural correlates of goal-directed and non-goal-directed movements. Proc. Natl. Acad. Sci. U. S. A. 118, 2021 (2021).

71. Fu, Z. et al. The geometry of domain-general performance monitoring in the human medial frontal cortex. Science (80-.). 376, (2022).

72. Cortes, C., Vapnik, V. & Saitta, L. Support-Vector Networks Editor. Mach. Leaming 20, 273–297 (1995).

73. Hennig, J. et al. How learning unfolds in the brain: toward an optimization view: Neuron. Neuron (2021).

74. Wang, J. X. et al. Prefrontal cortex as a meta-reinforcement learning system. Nat. Neurosci. 21, 860–868 (2018).

75. Curtis, C. E. & Lee, D. Beyond working memory: The role of persistent activity in decision making. Trends Cogn. Sci. 14, 216–222 (2010).

76. Tsujimoto, S., Genovesio, A. & Wise, S. P. Frontal pole cortex: Encoding ends at the end of the endbrain. Trends Cogn. Sci. 15, 169–176 (2011).

77. Compte, A., Brunel, N., Goldman-Rakic, P. S. & Wang, X. J. Synaptic Mechanisms and Network Dynamics Underlying Spatial Working Memory in a Cortical Network Model. Cereb. Cortex 10, 910–923 (2000).

78. Barbosa, J. et al. Interplay between persistent activity and activity-silent dynamics in the prefrontal cortex underlies serial biases in working memory. Nat. Neurosci. 2020 238 23, 1016–1024 (2020).

79. Koechlin, E. Prefrontal executive function and adaptive behavior in complex environments. Curr. Opin. Neurobiol. 37, 1–6 (2016).

80. Averbeck, B. & O’Doherty, J. P. Reinforcement-learning in fronto-striatal circuits. Neuropsychopharmacol. 2021 471 47, 147–162 (2021).

81. Lee, D., Seo, H. & Jung, M. W. Neural Basis of Reinforcement Learning and Decision Making. Annu. Rev Neurosc. 35, 287–308 (2012).

82. Lee, D. & Seo, H. Mechanisms of Reinforcement Learning and Decision Making in the Primate Dorsolateral Prefrontal Cortex. Ann. N. Y. Acad. Sci. 1104, 108–122 (2007).

83. Lim, DH., Yoon, Y.J., Her, E. et al. Active maintenance of eligibility trace in rodent prefrontal cortex. Sci Rep 10, *18860* (2020). doi: 10.1038/s41598-020-75820-0

84. Parker, N. F. et al. Choice-selective sequences dominate in cortical relative to thalamic inputs to NAc to support reinforcement learning. Cell Rep. 39, 110756 (2022).

85. Harvey, C. D., Coen, P. & Tank, D. W. Choice-specific sequences in parietal cortex during a virtual-navigation decision task. Nature (2012). doi:10.1038/nature10918

86. Koay, S. A., Charles, A. S., Thiberge, S. Y., Brody, C. D. & Tank, D. W. Sequential and efficient neural-population coding of complex task information. Neuron 110, 328–349.e11 (2022).

87. Asaad, F. W., Rainer, G. & Miller, E. K. Neural Activity in the Primate Prefrontal Cortex during Associative Learning. Neuron 21, 1399–1407 (1998).

88. Bichot, N. P., Schall, J. D. & Thompson, K. G. Visual feature selectivity in frontal eye fields induced by experience in mature macaques. Nature 381, 697–699 (1996).

89. White, I. M. & Wise, S. P. Rule-dependent neuronal activity in the prefrontal cortex. Exp. Brain Res. 126, 315–335 (1999).

90. Yang, G. R., Joglekar, M. R., Song, H. F., Newsome, W. T. & Wang, X. J. Task representations in neural networks trained to perform many cognitive tasks. Nat. Neurosci. 2019 222 22, 297–306 (2019).

91. Kirkpatrick, J. et al. Overcoming catastrophic forgetting in neural networks. doi:10.1073/pnas.1611835114

92. Sommer, M. A. & Wurtz, R. H. Visual perception and corollary discharge. Perception 37, (2008).

93. Sommer, M. A. & Wurtz, R. H. Brain circuits for the internal monitoring of movements. Annu. Rev. Neurosci. 31, 317 (2008).

94. Neupane, S., Guitton, D. & Pack, C. C. Two distinct types of remapping in primate cortical area V4. Nat. Commun. 7, 10402 (2016).

95. Duhamel, J. R., Colby, C. L. & Goldberg, M. E. The updating of the representation of visual space in parietal cortex by intended eye movements. Science 255, 90–2 (1992).

96. Marino, A. C. & Mazer, J. A. Perisaccadic Updating of Visual Representations and Attentional States: Linking Behavior and Neurophysiology. Frontiers (Boulder). 10, (2016).

97. Marino, A. C. & Mazer, J. A. Saccades Trigger Predictive Updating of Attentional Topography in Area V4. Neuron 98, 429–438.e4 (2018).

98. Mays, L. E. & Sparks, D. L. Dissociation of visual and saccade-related responses in superior colliculus neurons - PubMed. J. Neurophysiol. 43, 207–232 (1980).

99. Wurtz, R. H. Corollary Discharge Contributions to Perceptual Continuity Across Saccades. 10.1146/annurev-vision-102016-061207 4, 215–237 (2018).

100. Alexander, W. H. & Brown, J. W. Medial prefrontal cortex as an action-outcome predictor. Nat. Neurosci. 2011 1410 14, 1338–1344 (2011).

101. Heinzle, J., Aponte, E. A. & Stephan, K. E. Computational models of eye movements and their application to schizophrenia. Heinzle, Jakob; Aponte, Eduardo A; Stephan, Klaas E (2016). Comput. Model. eye movements their Appl. to Schizophr. Curr. Opin. Behav. Sci. 1121-29. 11, 21–29 (2016).

102. Adámek, P., Langová, V. & Horáček, J. Early-stage visual perception impairment in schizophrenia, bottom-up and back again. Schizophr. 2022 81 8, 1–12 (2022).

103. Broerse, A., Crawford, T. J. & Den Boer, J. A. Parsing cognition in schizophrenia using saccadic eye movements: a selective overview. Neuropsychologia 39, 742–756 (2001).

104. Egaña, J. I. et al. Small saccades and image complexity during free viewing of natural images in schizophrenia. Front. Psychiatry 4, 37 (2013).

105. Sutton, R. S., Barto, A. G. & Book, A. B. Reinforcement Learning: An Introduction. (2017).

106. Gnadt, J. W. & Andersen, R. A. Memory related motor planning activity in posterior parietal cortex of macaque. Exp. brain Res. 70, 216–220 (1988).

107. Ding, L. & Gold, J. I. Caudate encodes multiple computations for perceptual decisions. J. Neurosci. 30, 15747–15759 (2010).

108. Andersen, R. A., Snyder, L. H., Bradley, D. C. & Xing, J. Multimodal representation of space in the posterior parietal cortex and its use in planning movements. Annu. Rev. Neurosci. 20, 303–330 (1997).

109. Snyder, L. H. Coordinate transformations for eye and arm movements in the brain. Curr. Opin. Neurobiol. 10, 747–754 (2000).

110. Golomb, J. D., Chun, M. M. & Mazer, J. A. The native coordinate system of spatial attention is retinotopic. J. Neurosci. 28, 10654–10662 (2008).

111. Killian, N. J., Potter, S. M. & Buffalo, E. A. Saccade direction encoding in the primate entorhinal cortex during visual exploration. Proc. Natl. Acad. Sci. U. S. A. 112, 15743– 15748 (2015).

112. Meister, M. L. R. & Buffalo, E. A. Getting directions from the hippocampus: The neural connection between looking and memory. Neurobiol. Learn. Mem. 134 **Pt A**, 135–144 (2016).

113. Baram, A. B., Muller, T. H., Nili, H., Garvert, M. M. & Behrens, T. E. J. Entorhinal and ventromedial prefrontal cortices abstract and generalize the structure of reinforcement learning problems. Neuron 109, 713–723.e7 (2021).

114. Garvert, M. M., Dolan, R. J. & Behrens, T. E. J. A map of abstract relational knowledge in the human hippocampal–entorhinal cortex. Elife 6, (2017).

115. Krumin, M., Lee, J. J., Harris, K. D. & Carandini, M. Decision and navigation in mouse parietal cortex. Elife (2018). doi:10.7554/eLife.42583

116. Sailer, U., Flanagan, J. R. & Johansson, R. S. Eye–Hand Coordination during Learning of a Novel Visuomotor Task. J. Neurosci. 25, 8833–8842 (2005).

117. Land, M. F. Eye movements and the control of actions in everyday life. Prog. Retin. Eye Res. 25, 296–324 (2006).

118. Gerstner, W., Kempter, R., Van Hemmen, J. L. & Wagner, H. A neuronal learning rule for sub-millisecond temporal coding. Nat. 1996 3836595 383, 76–78 (1996).

119. Gilson, M., Burkitt, A. & Leo van Hemmen, J. STDP in recurrent neuronal networks. Front. Comput. Neurosci. 4, 23 (2010).

120. Gerstner, W., Lehmann, M., Liakoni, V., Corneil, D. & Brea, J. Eligibility Traces and Plasticity on Behavioral Time Scales: Experimental Support of NeoHebbian Three-Factor Learning Rules. Frontiers in Neural Circuits 12, 53 (2018).

121. Perrin, E. & Venance, L. Bridging the gap between striatal plasticity and learning. Current Opinion in Neurobiology 54, 104–112 (2019).

122. Bittner, K. C., Milstein, A. D., Grienberger, C., Romani, S. & Magee, J. C. Behavioral time scale synaptic plasticity underlies CA1 place fields. Science (80-.). 357, 1033–1036 (2017).

123. Suvrathan, A. Beyond STDP — towards diverse and functionally relevant plasticity rules. Current Opinion in Neurobiology 54, 12–19 (2019).

124. Mansvelder, H., Verhoog, M. & Goriounova, N. Synaptic plasticity in human cortical circuits: cellular mechanisms of learning and memory in the human brain? Curr. Opin. Neurobiol. 54, 186–193 (2019).

125. Stokes, M. G. ‘Activity-silent’ working memory in prefrontal cortex: A dynamic coding framework. Trends in Cognitive Sciences 19, 394–405 (2015).

126. Mongillo, G., Barak, O. & Tsodyks, M. SynaptiC Theory of Working Memory. Science *(80-.).* **319**, 1543–1546 (2008).

## References

[1] Evarts, E. V. Pyramidal tract activity associated with a conditioned hand movement in the monkey. J. Neurophysiol. 29, 1011–1027 (1966).

[2] Judge, S. J., Richmond, B. J. and Chu, F. C. Implantation of magnetic search coils for measurement of eye position: an improved method. Vision Res. 20, 535–538 (1980).

[3] Suner, S., Fellows, M. R., Vargas-Irwin, C., Nakata, G. K. and Donoghue, J. P. Reliability of Signals From a Chronically Implanted, Silicon-Based Electrode Array in Non-Human Primate Primary Motor Cortex. IEEE Trans. Neural Syst. Rehabil. Eng. 13, 524–541 (2005).

[4] Santhanam, G., Ryu, S. I., Yu, B. M., Afshar, A. and Shenoy, K. V. A high­-performance brain-computer interface. Nature 442, 195–198 (2006).

[5] Mould, M. S., Foster, D. H., Amano, K. and Oakley, J. P. A simple nonparametric method for classifying eye fixations. Vision Res. 57, 18–25 (2012).

[6] Bruce, C. J. and Goldberg, M. E. Primate frontal eye fields. I. Single neurons discharging before saccades. J. Neurophysiol. 53, 603–35 (1985).

[7] A Graf, A. B., Kohn, A., Jazayeri, M. and Anthony Movshon, J. Decoding the activity of neuronal populations in macaque primary visual cortex. Nat. Neurosci. 14, (2011).

[8] DeAngelis, G. C., Ohzawa, I. and Freeman, R. D. Receptive-field dynamics in the central visual pathways. Trends Neurosci. 18, 451–458 (1995).

